# Long-Range Input to Cortical Microcircuits Shapes EEG-BOLD Correlation

**DOI:** 10.1101/2025.06.06.658058

**Authors:** Vincent S.C. Chien, Stanislav Jiříček, Thomas R. Knösche, Jaroslav Hlinka, Helmut Schmidt

## Abstract

Electroencephalography (EEG) rhythms and blood-oxygen-level-dependent (BOLD) activity, though generated by different mechanisms, exhibit correlations. The level of correlation varies between EEG frequency bands, brain regions, and experimental paradigms, but the underpinning mechanisms of this correlation remain poorly understood. Here we create a mathematical, data-informed model of a cortical microcircuit that encompasses all major neuron types across cortical layers, and use it to generate EEG and BOLD under various external input conditions. The model exhibits noise-driven fluctuations giving rise to distinct EEG rhythms, with external inputs modulating EEG spectral characteristics. In line with experimental findings, we observe negative alpha-BOLD correlations and positive gamma-BOLD correlations across different input configurations. Temporal variability of the input is found to increase EEG-BOLD correlation and to improve the correspondence with experimental results. This study provides a mathematical framework to theoretically study the correlation of EEG and BOLD features in a comprehensive way.

## Introduction

Electroencephalography (EEG) and functional magnetic resonance imaging (fMRI) are widely used non-invasive methods for imaging the human brain. EEG records the electric potentials on the scalp that reflect primarily the synaptic activity at the pyramidal cells (Buzsáki et al., 2012), whereas fMRI measures the blood-oxygen-level-dependent (BOLD) signal that reflects the metabolism of synaptic activity (Howarth et al., 2021). The two modalities provide complementary information for inferring the underlying cortical neural activity. Combining EEG and fMRI thus offers valuable insights into their neural origins.

The alpha rhythm is a prominent feature in human EEG over occipital areas first observed by Berger in 1929 (Berger, 1929). Functionally, alpha activity has been associated with task-negative states, exhibiting higher levels during rest compared to task performance (Clayton et al., 2018; Magosso et al., 2019). Simultaneous EEG-fMRI studies have found that alpha power is negatively correlated with BOLD signals in occipital regions and positively correlated with BOLD signals in areas such as the insula and thalamus (Goldman et al., 2002; Laufs, Kleinschmidt, et al., 2003; Laufs, Krakow, et al., 2003; Moosmann et al., 2003). At an individual level, the alpha-BOLD correlation exhibits intra-subject and inter-subject variability (Gonçalves et al., 2006). The gamma rhythm, usually anticorrelated with the alpha and beta rhythms, has been associated with sensory processing and tasks such as working memory (Shu et al., 2024). Local field potential (LFP) recordings, as opposed to spatially coarse-grained EEG recordings, can record this spatially localised, high-frequency neural activity at high signal-to-noise ratios. A simultaneous LFP-fMRI study demonstrated that gamma power correlates positively with BOLD responses and visual stimulus intensity (Niessing et al., 2005). In short, simultaneous EEG-fMRI and LFP-fMRI recordings are helpful for investigating the location of brain rhythms and their relationship with the BOLD signal.

A few theoretical studies have sought to provide mechanistic explanations that integrate these experimental observations, but many questions remain unresolved. For instance, while alpha rhythms are generally attributed to network-level mechanisms (Halgren et al., 2019), their precise origin and relationship with BOLD activity are still debated. It was demonstrated that source-localised alpha power predicts BOLD activity and reproduces related phenomena in a whole-brain modeling framework (Schirner et al., 2018). However, it was assumed that a causal link exists between alpha power and BOLD without addressing the origin of alpha rhythms. By contrast, a thalamocortical model exhibited anticorrelations between the alpha power of the simulated EEG and a low-frequency BOLD proxy (Pang & Robinson, 2018), where the alpha rhythms were attributed to delays in the thalamocortical loop and explained alpha-BOLD anticorrelations by variations in corticothalamic and intrathalamic feedback strengths. However, this model did not address fast time-varying effects such as stimulus intensity on internal brain dynamics. Thus, a more comprehensive theoretical framework that integrates the generation of EEG rhythms and their relationships with BOLD signals is still lacking.

Given that both EEG and BOLD signals predominantly originate from postsynaptic potentials (PSPs) (Biasiucci et al., 2019; Ekstrom, 2010), and considering that distinct inhibitory neuron types influence EEG rhythms differently (Whittington & Traub, 2003) and exhibit varied neurovascular coupling properties (Cauli et al., 2004), a cortical microcircuit model incorporating multiple inhibitory neuron types offers a promising framework for investigating local neural dynamics underlying EEG rhythms and their relationship with BOLD signals. Such a mesoscopic model has the advantage of providing insights into interactions between neural populations and potential factors contributing to heterogeneity in EEG-BOLD correlations, as observed in our experimental findings (Figure 1A-D).

**Figure 1.**
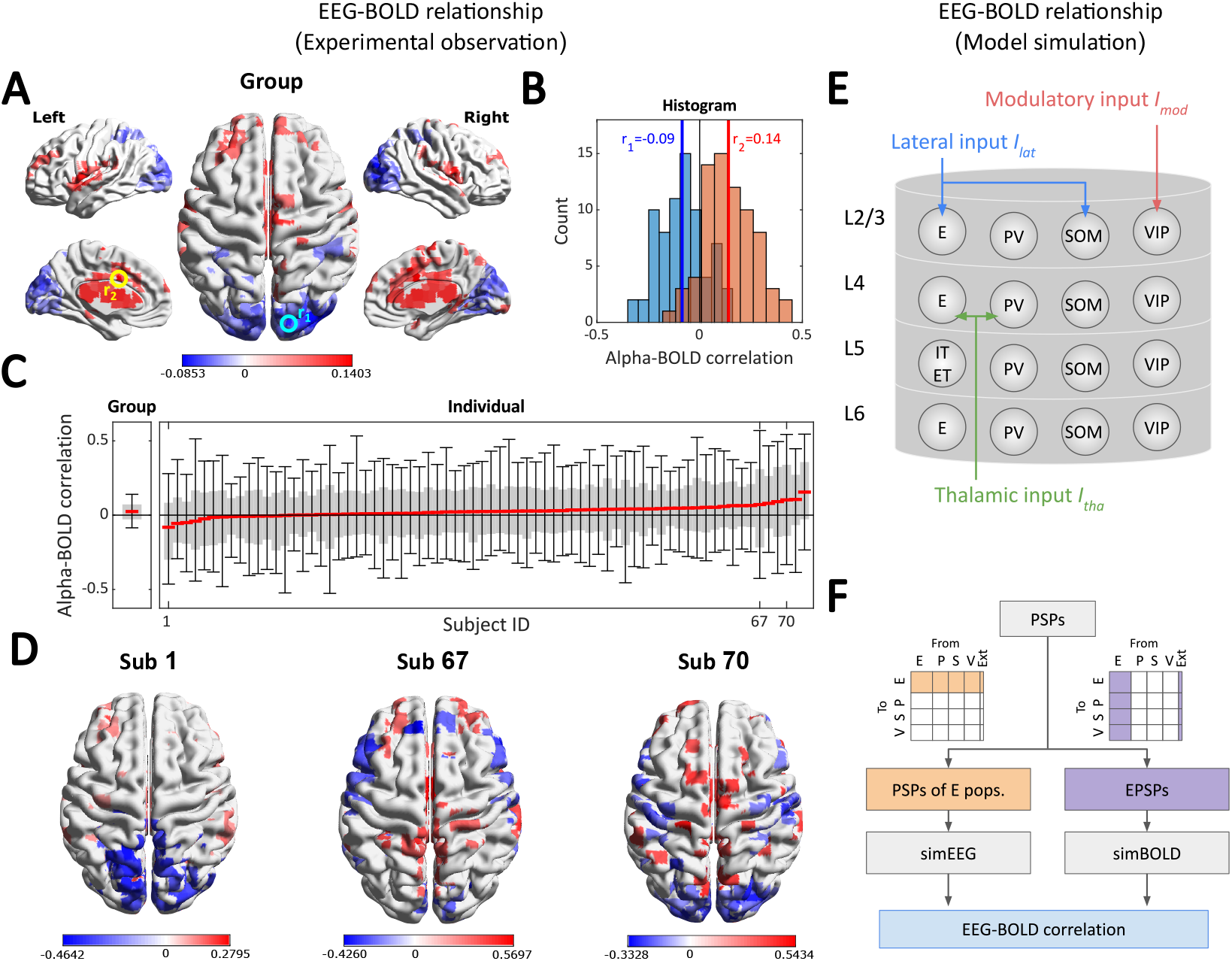
Heterogeneity of alpha-BOLD correlation and modeling approach. (**A**) Group-level alpha-BOLD correlation map across 72 subjects. Values from 5% to 95% are masked. (**B**) Histograms of alpha-BOLD correlations at two example voxels, highlighted by the yellow (middle cingulate gyrus) and cyan (middle occipital gyrus) circles in (**A**). Box plot of group-level (averaged across subjects) and individual, voxel-wise alpha-BOLD correlations. Subjects are sorted by their median correlation values (red lines). Gray boxes represent the 5th to 95th percentiles, and whiskers indicate the minimum and maximum alpha-BOLD correlations. (**D**) Three individual examples: Subject 1 has the most negative median correlation, Subject 67 displays the widest range of correlations, and Subject 70 exhibits the highest spatial similarity with the group-level correlation map. (**E**) Model setup. The cortical microcircuit model consists of 17 neural populations, including excitatory (E, IT, and ET) and inhibitory (PV, SOM, and VIP) populations across cortical layers L2/3, L4, L5, and L6. The model receives three types of external inputs: lateral (*I*_lat_), modulatory (*I*_mod_), and thalamic (*I*_tha_). Internal connections are not shown for clarity. (**F**) Model simulation. The simulated EEG signal (simEEG) is generated from the postsynaptic potentials (PSPs) at all excitatory populations (orange section), and the simulated BOLD signal (simBOLD) is generated from the PSPs by all excitatory inputs (purple section). EEG-BOLD correlation calculations follow experimental protocols (see Methods for details).

While the neurovascular coupling process has been explored extensively in previous studies (Cauli et al., 2004; Howarth et al., 2021), our primary focus here is not to provide a detailed biophysical account of hemodynamic mechanisms. Instead, we aim to clarify how intrinsic neural dynamics, particularly the interplay between excitatory and multiple inhibitory cell types, shape the observed EEG-BOLD correlations. By emphasizing neural population interactions, rather than vascular biophysics, the model highlights how changes in long-range input configurations can lead to diverse EEG-BOLD relationships across cortical regions. We developed a cortical microcircuit model that incorporates excitatory (E) and inhibitory (PV, SOM, and VIP) populations across cortical layers (L2/3, L4, L5, and L6) with realistic configurations. The model receives three types of external inputs: (1) lateral input from neighboring cortical microcircuits, targeting pyramidal cells and SOM interneurons in L2/3 (Adesnik et al., 2012; Yavorska & Wehr, 2016); (2) modulatory input from higher order cortical areas targeting VIP interneurons in L2/3 (Hertäg & Sprekeler, 2019); and (3) thalamic input targeting pyramidal cells and PV interneurons in L4 (Kloc & Maffei, 2014). In the simulations, Gaussian noise is independently applied to all synaptic connections, and EEG and BOLD signals are derived from the

PSPs (Figure 1F). The analysis begins with the model behavior where noise-driven fluctuations resembling EEG rhythms are observed. The effect of synaptic noise and input variability on EEG-BOLD temporal correlations is then explored, with a simulation example replicating the LFP-BOLD correlations reported in (Magri et al., 2012). Lastly, the contributions of individual neural populations are examined to investigate the sources of EEG rhythms and EEG-BOLD correlations.

The focus on neural dynamics provides a mechanistic bridge between mesoscopic circuit activity and macroscopic EEG-fMRI observations. It enables us to explore how transitions between network states, mediated by different input pathways, modulate both spectral power and BOLD amplitude. The framework thus serves as a tool to investigate experimental findings such as the heterogeneity of EEG-BOLD correlations and their dependence on brain states.

## Results

### Heterogeneity of alpha-BOLD correlation

In Figure 1A, we show a group-level alpha-BOLD correlation map across 72 healthy participants. Negative alpha-BOLD correlations are found in the parietal and occipital areas, with a minimum in a voxel in the middle occipital gyrus (*r*_1_, highlighted by the cyan circle). Positive alpha-BOLD correlations are distributed in frontal and medial brain areas, with a maximum in a voxel in the middle cingulate gyrus (*r*_2_, highlighted by the yellow circle). We select these two voxels and show the histograms across subjects in Figure 1B. The variability in alpha-BOLD correlation across individual subjects is substantial (*r*_1_ = −0.09 ± 0.11, and *r*_2_ = 0.14 ± 0.12), aligning with findings from a previous study (Gonçalves et al., 2006). In Figure 1C, we present the brain-wide distribution of individual and group-level alpha-BOLD correlations. The variability across brain voxels in individual alpha-BOLD correlations is also notably greater than that observed in the group-level correlations. Figure 1D shows the alpha-BOLD correlation maps of three representative participants to further illustrate the heterogeneity of alpha-BOLD correlation maps. The heterogeneity is evident at multiple levels (i.e., across brain voxels and between subjects). We attribute this heterogeneity to the noise in synaptic activity and the variability in long-range input to cortical microcircuits, as demonstrated later through simulations.

### Noise-driven fluctuations

The external inputs to the cortical microcircuit model — lateral input *I*_lat_, modulatory input *I*_mod_, and thalamic input *I*_tha_ — determine whether it operates in an oscillatory or steady state. In this study, we focus on the range of external inputs where the model remains in a steady state without oscillatory behavior, and where it therefore exhibits noise-driven fluctuations when independent Gaussian noise is applied to internal connections.

In Figure 2, we characterise the spectral properties of the noise-driven fluctuations under constant input conditions (input range *I*_lat_ = [2.5–4.5], *I*_mod_ = [0–25], and *I*_tha_ = [0–5]). The upper limits of these intervals ensure that the system remains in the steady state. The lower limit of *I*_lat_ = 2.5 is motivated by the fact that for smaller values, the system displays relatively uniform behavior with little variability in the EEG spectra. Furthermore, cortical microcircuits receive continuous input from nearby areas due to prevalent lateral connections (Grinvald et al., 1994), which would justify the non-zero lower limit of *I*_lat_. Figure 2A illustrates the power change in decibel (dB) in frequency bands delta (0.5– 4 Hz), theta (4–7 Hz), alpha (8–12 Hz), beta (13–30 Hz), and gamma (30–50 Hz) relative to baseline power at *I*_lat_, *I*_mod_, and *I*_tha_ = 0. Overall, the modulatory input *I*_mod_ induces a distinct switching behavior, altering the role of thalamic input between sharply contrasting regimes. At lower *I*_mod_, the dominant frequency power of noise-driven fluctuations transitions from alpha to beta and gamma as thalamic input *I*_tha_ increases. At higher *I*_mod_, substantial delta and theta power emerge. This switching behavior becomes more pronounced with increasing lateral input *I*_lat_, characterised by enhanced band power and a shifted dependence on thalamic input, such as the observed shift in theta power within the input space.

**Figure 2.**
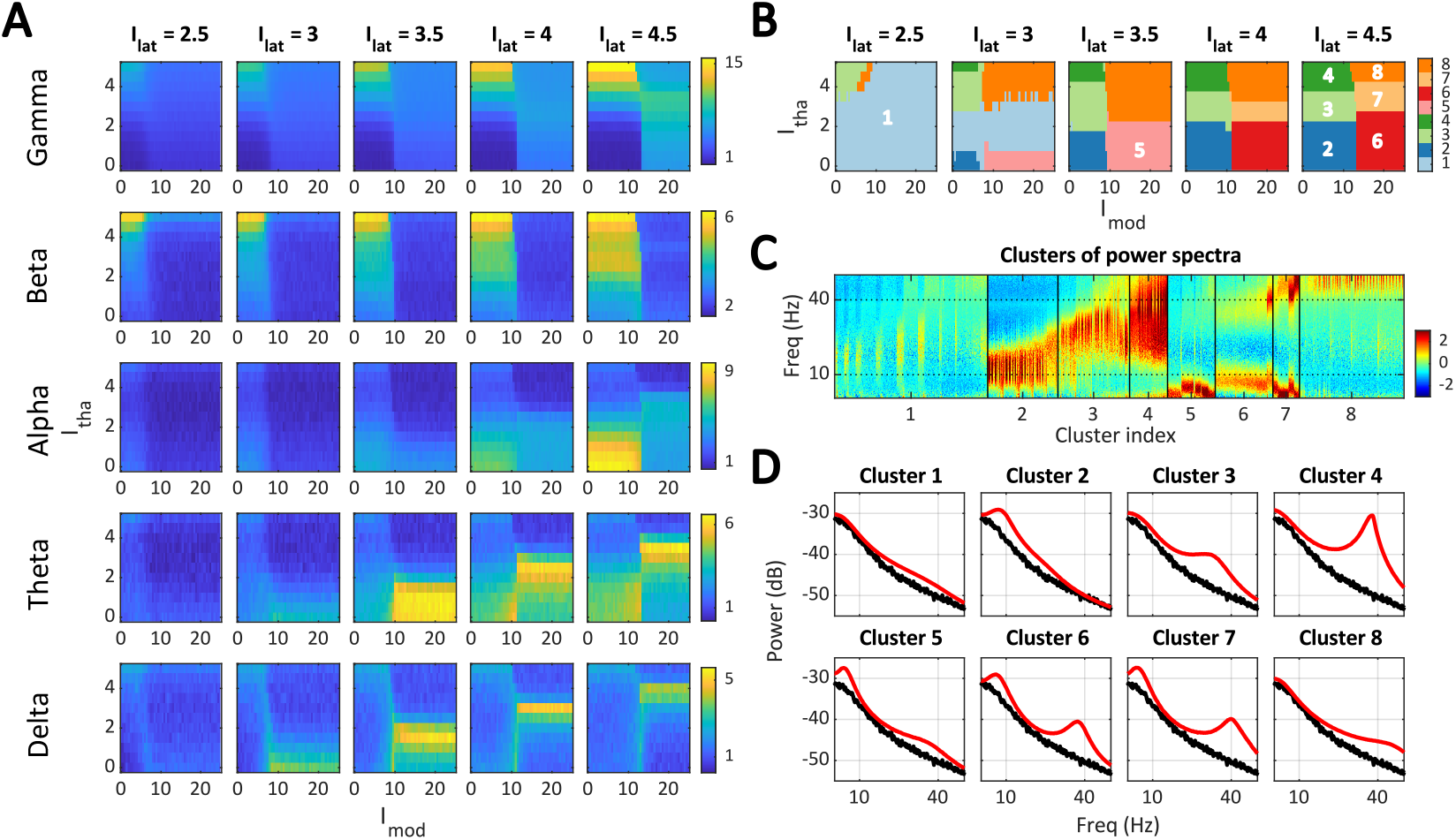
Spectral properties of noise-driven fluctuations. (**A**) Relative EEG power (in dB) compared to the baseline condition (*I*_lat_, *I*_mod_, *I*_tha_ = 0) across frequency bands: gamma (30–50 Hz), beta (13–30 Hz), alpha (8–12 Hz), theta (4–7 Hz), and delta (0.5–4 Hz), shown from top to bottom. (**B**) Eight K-means clusters by z-scored EEG spectral features (0.5–50Hz). (**C**) The z-scored power spectra were sorted by cluster index to visually inspect spectral heterogeneity within each cluster. (**D**) The mean spectra (red) for each cluster, alongside the baseline spectrum (black).

These spectral features align with experimental observations, such as EEG alpha in the eyes-closed resting state, corresponding to low *I*_tha_ (Barry et al., 2007), EEG gamma during sensory tasks, corresponding to high *I*_tha_ (Muthukumaraswamy & Singh, 2013), and EEG theta during attentional processes with concurrent reduction of alpha power, corresponding to high *I*_mod_ (Cona et al., 2020; Keller et al., 2017; Missonnier et al., 2006).

To facilitate visualization and conceptual understanding, we divide the input space into eight clusters based on the EEG power spectra (0.5–50 Hz) using K-means clustering (Figure 2BCD). The input space at low *I*_lat_ is dominated by Cluster 1 characterised by only small deviations of their spectra from the baseline spectra, which supports our choice of lower limit for *I*_lat_. In Clusters 2 to 4, where the modulatory input *I*_mod_ is low, fluctuations predominantly occur in the alpha, beta, and gamma bands. In Clusters 5 to 8, where the modulatory input *I*_mod_ is high, fluctuations predominantly occur in the delta, theta, and gamma bands. Note that the specific number of clusters is not critical but has emerged as the optimal number of clusters based on centroid distance (supplementary Figure S1).

### EEG alpha, gamma, and BOLD

We examine the BOLD amplitude and its relationship with EEG power in the alpha and gamma bands under constant input conditions. Figure 3A shows the mean BOLD amplitudes in the input space. Generally, BOLD amplitude is higher at increased levels of external inputs and reaches its peak in Cluster 4, where strong beta and gamma activity are observed. Figure 3B illustrates the relationship between EEG alpha power, gamma power, and BOLD amplitude. Gamma power generally shows a positive correlation with BOLD amplitude (with some exceptions in Cluster 8), whereas alpha power typically exhibits a negative correlation with BOLD amplitude (with exceptions in Clusters 4 and 6). This result offers an intuitive explanation for experimental observations of positive gamma-BOLD correlations (Logothetis et al., 2001; Magri et al., 2012) and negative alpha-BOLD correlations (Goldman et al., 2002; Laufs, Kleinschmidt, et al., 2003; Laufs, Krakow, et al., 2003; Moosmann et al., 2003). Here, the BOLD amplitude and EEG power are time-averaged quantities. It is still necessary to obtain the EEG-BOLD temporal correlation to investigate the experimental findings closely.

**Figure 3.**
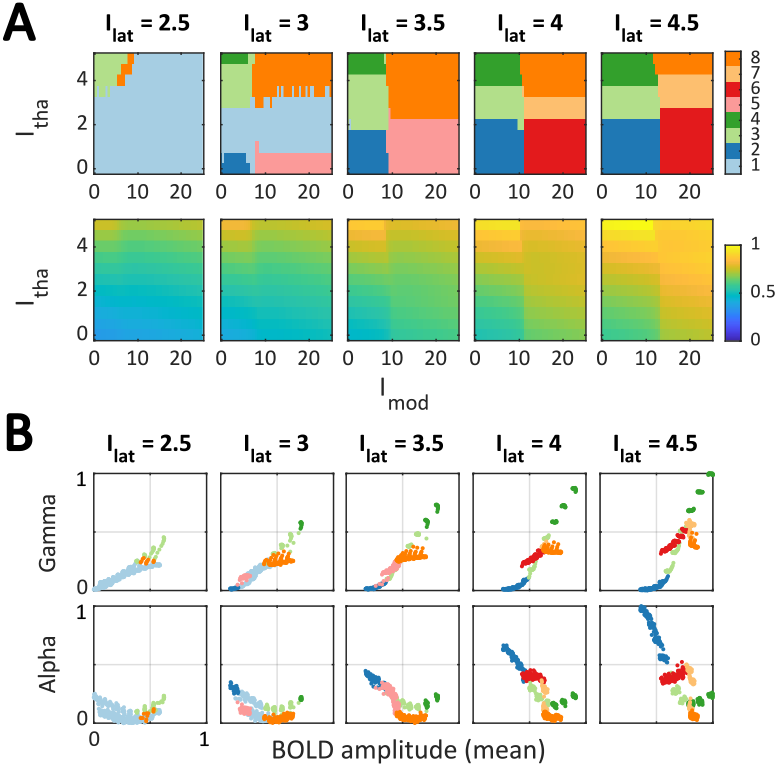
Mean BOLD amplitudes and corresponding EEG alpha and gamma powers. (**A**) K-means clusters (top row, same as in Figure 2**B**) and the mean BOLD amplitude (bottom row). (**B**) Scatter plots of gamma-BOLD (top) and alpha-BOLD (bottom) relationships. Mean BOLD amplitude (x-axis) and EEG band powers (y-axis) are normalised. Full version in supplementary Figure S2.

### EEG-BOLD temporal correlation: constant input condition

The constant input conditions involved simulating 10 minutes of simultaneous EEG and BOLD data 100 times for each combination of input levels. Figure 4A and B present the mean alpha-BOLD and mean gamma-BOLD correlations throughout the input space, respectively. We observe both positive and negative alpha-BOLD correlations, whereas the gamma-BOLD correlations are predominantly positive. Interestingly, the region of negative alpha-BOLD correlations does not align with the high alpha region (Cluster 2) but with a region where both alpha and gamma coexist (Cluster 6). Moreover, the region of positive alpha-BOLD correlations aligns with Cluster 7 where delta, theta, and gamma are strong (cf. Figure 2A). In the same region of input space, we observe large amplitude fluctuations of the BOLD signal, as quantified by the standard deviation of the time series (supplementary Figure S2C).

**Figure 4.**
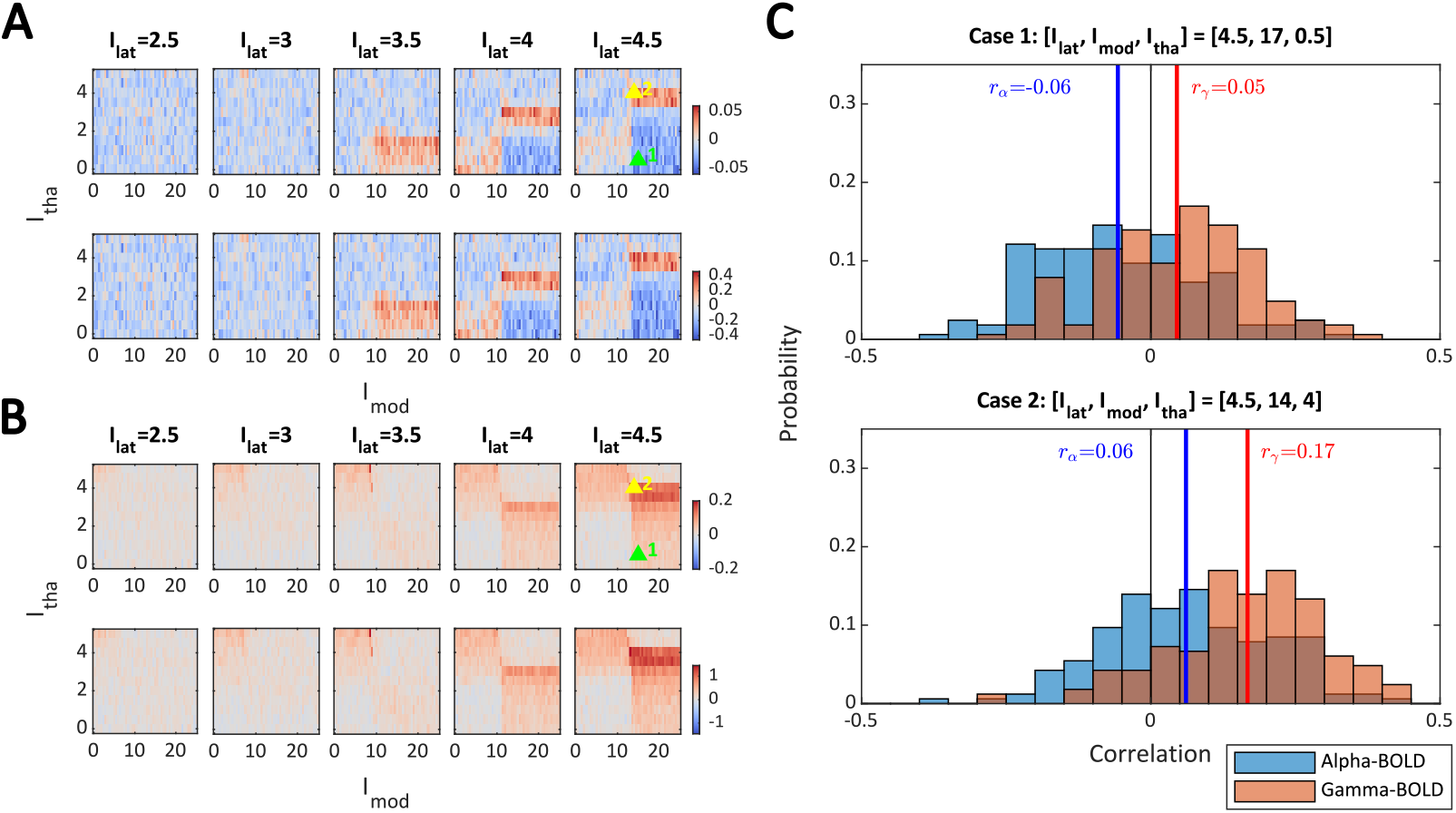
EEG-BOLD correlation under constant input conditions. (**A**) Mean alpha-BOLD correlation, and (**B**) mean gamma-BOLD correlation, across 100 runs (top row) and corresponding effect size (mean/std, bottom row). The correlations are calculated from *t* = [200–550] sec in each run. (**C**) Histogram for two example cases. In Case 1, the mean alpha-BOLD correlation *r*_*α*_ is negative (blue line), and the mean gamma-BOLD correlation *r*_*γ*_ is positive (red line). In Case 2, both *r*_*α*_ and *r*_*γ*_ are positive. The two example cases are indicated by green (case 1) and yellow (case 2) triangles in **A** and **B**.

Figure 4C presents two examples of the distribution of EEG-BOLD correlations across 100 runs. The mean EEG-BOLD correlations under constant input conditions are relatively low, around ± 0.05. The results remain stable even when the EEG dipole orientation and amplitude are randomised (see supplementary Figure S3), indicating that the observed EEG-BOLD correlations are minimally influenced by the specifics of the EEG forward model. Interestingly, the mean and range of simulated alpha-BOLD correlations in Figure 4C qualitatively correspond to distributions of empirical alpha-BOLD correlations of the most negatively/positively correlating voxels across subjects (Figure 1B) as well as whole brain alpha-BOLD correlations distributions for single subject maps (Figure 1C).

### EEG-BOLD temporal correlation: varying input condition

In the previous section, we demonstrated that synaptic noise can contribute to weak EEG-BOLD correlations. Here, we introduce variability in the external input, which more closely mimics realistic conditions. We illustrate three example cases of varying input conditions (Figure 5A): Case 1 (*I*_lat_ = 4.5, *I*_mod_ = [0–2], and *I*_tha_ = [0–2]) lies in the region of Cluster 2 where there is strong alpha power, Case 2 (*I*_lat_ = 4.5, *I*_mod_ = [12.5–14.5], and *I*_tha_ = [0–2]) spans Clusters 2 and 6, and Case 3 (*I*_lat_ = 4.5, *I*_mod_ = [16–18], and *I*_tha_ = [0–2]) lies in the region of Cluster 6. Figure 5B shows a single run of the EEG spectrogram, and the time series of alpha, gamma power, and BOLD amplitude for the three cases. In Cases 1 and 2, we observe negative alpha-BOLD correlations. Case 2 exhibits switches between two spectral patterns where alpha and gamma powers are anti-correlated. In Case 3, we observe a positive alpha-BOLD correlation, which would have been negative under constant input conditions. Figure 5C shows the histogram of alpha-BOLD and gamma-BOLD correlations across 100 runs. The mean Gamma-BOLD correlations are positive in all three cases, while the mean alpha-BOLD correlations vary substantially (from −0.66 in Case 1 to 0.16 in Case 3). The result shows that the effect of external input variability on EEG-BOLD correlations can be much greater than that of synaptic noise.

**Figure 5.**
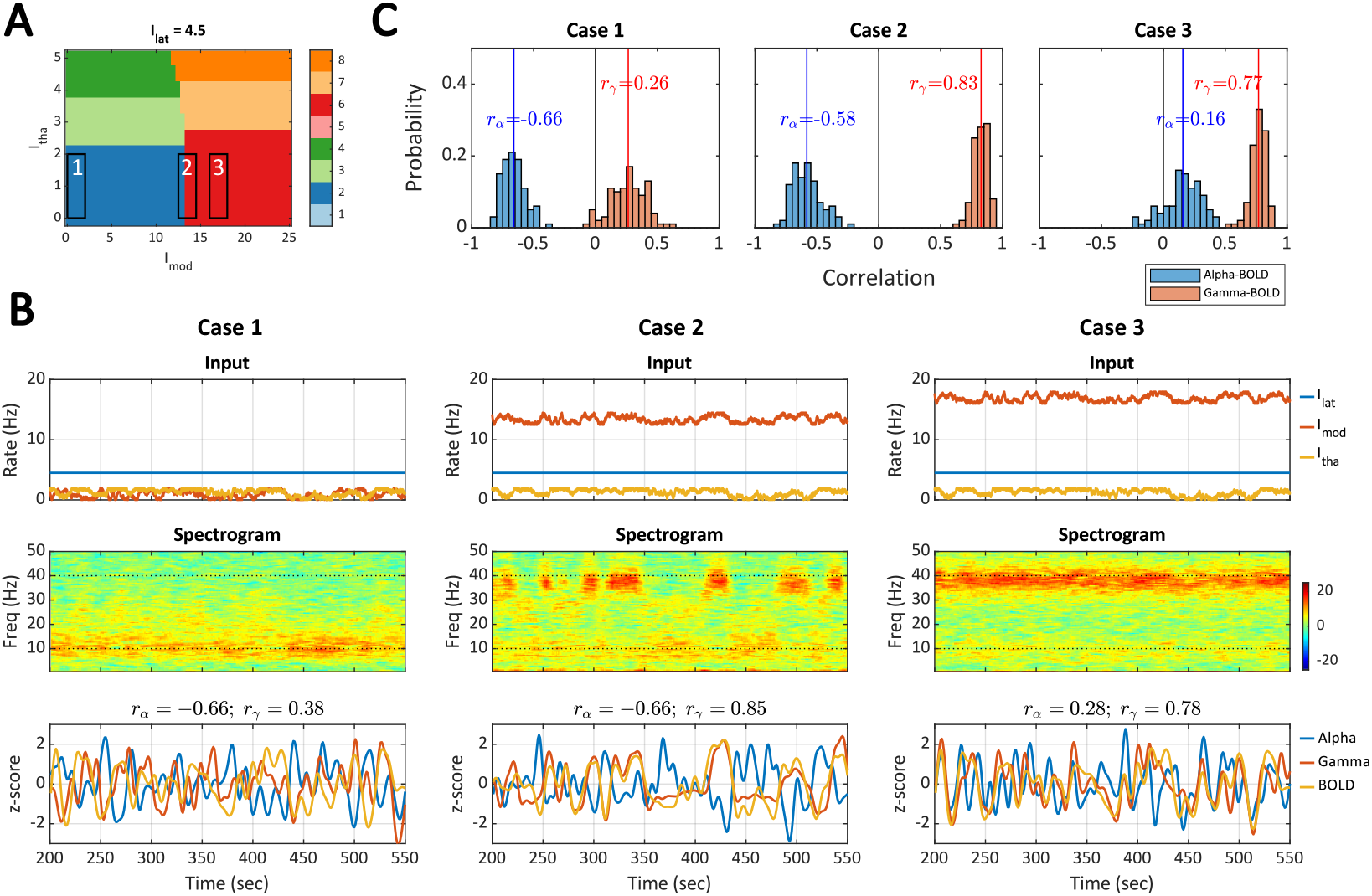
EEG-BOLD correlation under varying input conditions. (**A**) The K-means clusters (same as in Figure 2**B**) and three varying input cases where the input ranges are confined in the black rectangles: [*I*_lat_, *I*_mod_, *I*_tha_] = [4.5, 0–2, 0–2] (Case 1), [4.5, 12.5–14.5, 0–2] (Case 2), and [4.5, 16–18, 0–2] (Case 3). (**B**) A single simulation run. The top row shows the random walk inputs. The middle row shows the spectrogram of the simulated EEG signal. The bottom row shows the alpha power signal, the gamma power signal, and the simulated BOLD signal. The alpha-BOLD correlation *r*_*α*_ and gamma-BOLD correlation *r*_*γ*_ are calculated from *t* = [200–550] sec. (**C**) Histogram of alpha-BOLD (blue) and gamma-BOLD (orange) correlations across 100 runs.

The distinct alpha-BOLD correlations in the three example cases can be intuitively explained by viewing the distribution of mean alpha power and mean BOLD amplitude (second row in Figure 3B). In Case 1, the input ranges stay within Cluster 2 (blue dots), where the distribution of mean alpha power and mean BOLD amplitude forms a negative correlation. In Case 3, the input ranges stay within Cluster 6 (red dots), where the distribution of mean alpha power and mean BOLD amplitude forms a positive correlation. In Case 2, the input spans Clusters 2 and 6, resulting in an overall negative correlation.

Resonance to internal synaptic noise produces weak EEG-BOLD correlations (Figure 4), and introducing variability in external inputs can significantly modify these correlations (Figure 5). These two driving mechanisms shape the spectral properties of the cortical microcircuit model, resulting in diverse and dynamic EEG-BOLD relationships. The presence of variation in input amplitude enhances EEG-BOLD correlations and increases synchrony between neural populations. Supplementary Figures S8 and S9 provide examples demonstrating how greater variability in external inputs strengthens the correlation of neural firing activity.

### Replication of LFP-BOLD correlation

Alpha-BOLD anticorrelation and gamma-BOLD correlation have also been reported in localised brain regions in experimental studies. For instance, a simultaneous LFP-fMRI study reported alpha and gamma relationships with BOLD signals during spontaneous activity in the early visual cortices of anaesthetized monkeys (Magri et al., 2012). We validate our model by replicating their finding, where the *overall* alpha-BOLD correlation is positive, and the *fixed-power* alpha-BOLD correlation is negative (as shown in Figure 7 in Magri et al., 2012). We follow the methods described in (Magri et al., 2012). For overall correlation, we compute the Pearson correlation between LFP power at each frequency bin in the spectrogram (delayed by 3.5 seconds) and the BOLD signal. For fixed-power correlation, we divide the LFP power (the sum of the spectrogram from 0.5 to 100 Hz) into 10 percentile levels and compute the Pearson correlation at each level. Note that in this study, we use “LFP” and “EEG” inter-changeably.

We simulate 10 minutes of LFP and BOLD using random, time-varying inputs for 100 runs. Figure 6A shows a single run of simulation, with inputs confined to *I*_lat_ = [2.5–3], *I*_mod_ = [0–25], and *I*_tha_ = [0–5]. The external inputs traverse Clusters 1, 2, 3, 4, 5, and 8 (indicated by colored stripes beneath the input sub-plot), and the spectrogram reflects corresponding spectral transitions. The LFP power and the BOLD signal are highly correlated. Figure 6B shows 2D histograms and correlations among LFP power, alpha power, gamma power, and BOLD amplitude in 100 runs. The overall gamma-BOLD, gamma-LFP, and LFP-BOLD correlations are positive. Figure 6C shows the overall correlation (left) and the fixed-power correlation (middle) for each frequency bin. The overall correlations are positive across frequencies (red curve), whereas the mean fixed-power correlations can be negative at lower frequencies (black curve). Figure 6D shows that the fixed-power alpha-BOLD correlations for each percentile level are predominantly negative. These locally negative alpha-BOLD distributions (magenta ellipses) combine and produce a globally positive alpha-BOLD distribution (white ellipse), providing an intuitive interpretation of Figure 6C and the experimental findings in Figure 7 in (Magri et al., 2012). Figure 6E details the composition of spectral clusters at each percentile level for additional information. The external inputs traveling in Clusters 3, 4, and 8 (corresponding to a higher thalamic input *I*_tha_) contribute to the higher LFP power percentiles, whereas the external inputs traveling in Clusters 1, 2, and 5 (corresponding to lower *I*_tha_) contribute to the lower LFP power percentiles.

**Figure 6.**
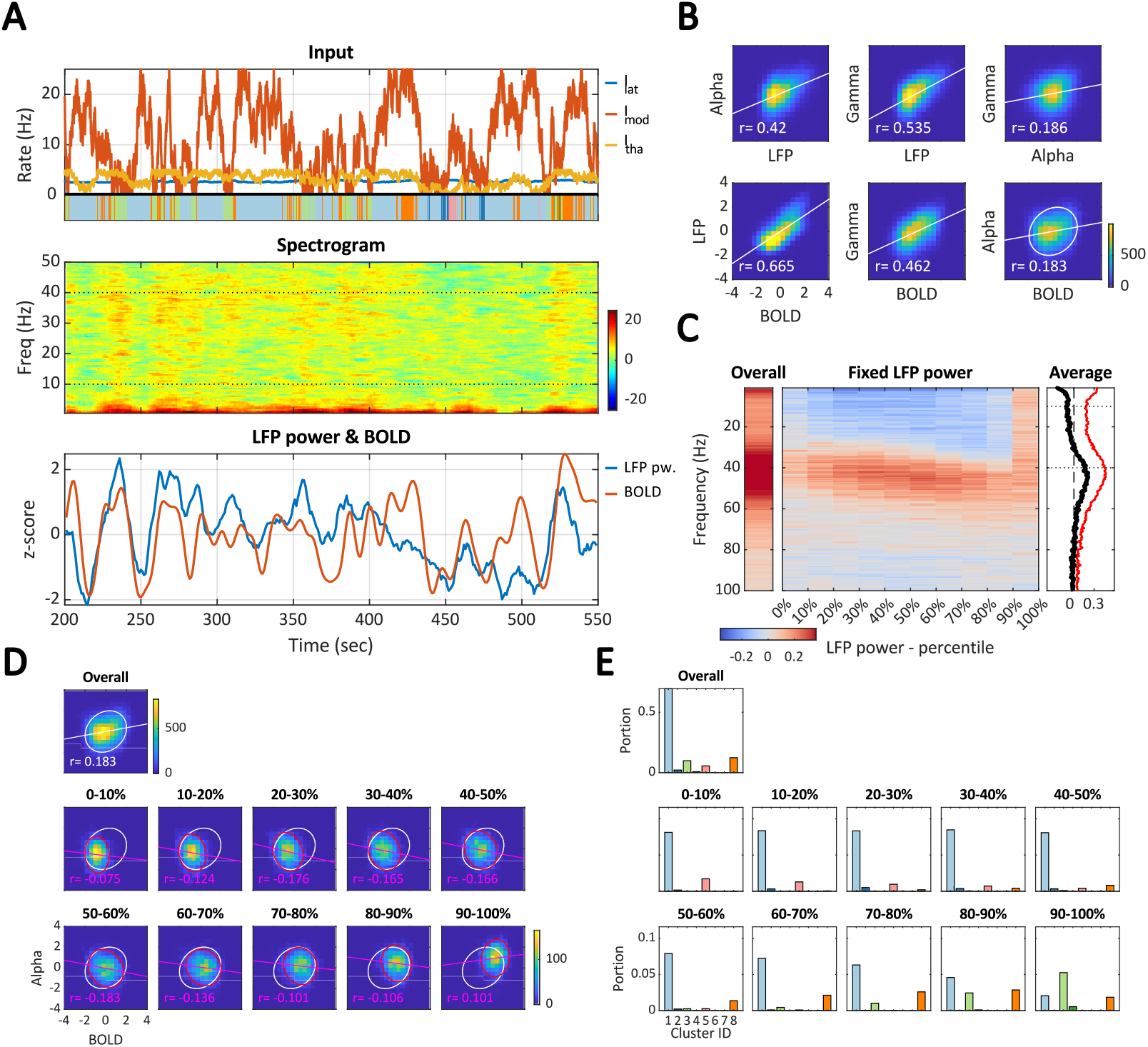
Reproducing the LFP-BOLD correlation from (Magri et al., 2012). (**A**) A single 10-minute simulation (start and end segments not shown). Top row: Varying inputs are constrained within *I*_lat_ = [2.5–3], *I*_mod_ = [0–25], and *I*_tha_ = [0–5]. Colored stripes beneath indicate the corresponding K-means cluster IDs. Middle row: LFP spectrogram (only showing 0.5–50 Hz for visual clarity). Bottom row: Time series of overall LFP power (sum of spectrogram from 0.5–100 Hz) and BOLD amplitude. (**B**) 2D histogram for overall LFP power, alpha power, gamma power, and BOLD amplitude across 100 runs. (**C**) Overall LFP-BOLD correlation at individual frequencies, averaged over 100 runs (left); fixed-power LFP-BOLD correlation, where the LFP power is divided into 10 percentile levels (middle); and average fixed-power correlation (black curve) compared with the overall correlation (red curve). (**D**) 2D histograms showing the alpha-BOLD distribution for overall correlation (with a white ellipse) and fixed-power correlations (with magenta ellipses) (**E**) Composition of spectral clusters for each power percentile level (cluster ID color code as in Figure 2B).

**Figure 7.**
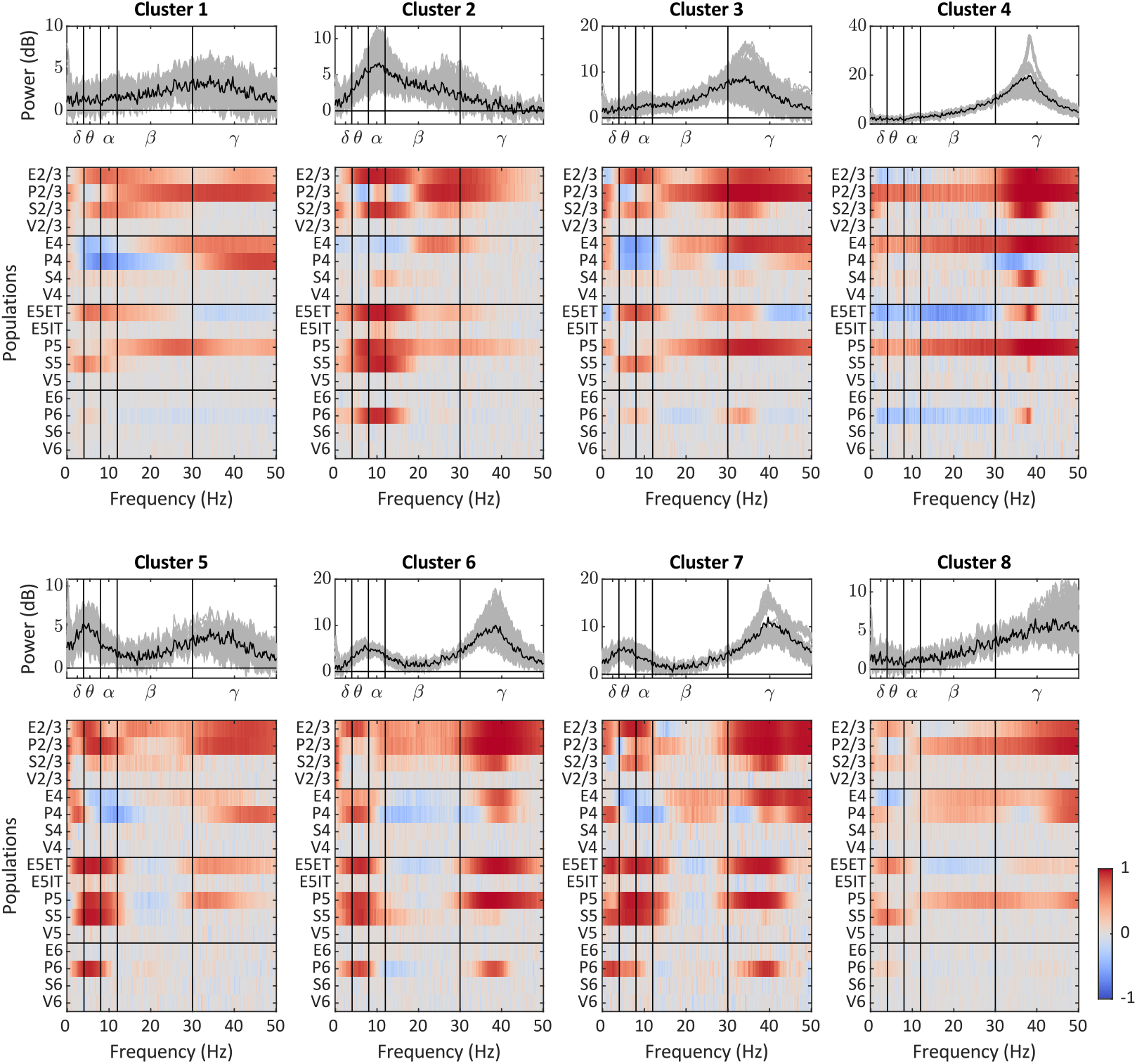
Relationship between the spectra of EEG and PSPs. The first and third rows display the baseline-corrected EEG power spectra (spectEEG) for k=8 clusters within the input space *I*_lat_ = [2.5–4.5], *I*_mod_ = [0–25], and *I*_tha_ = [0–5]. The second and fourth rows show the similarity (measured by correlation over input space) between spectEEG and spectPSP, where spectPSP (shown in supplementary Figure S7C) represents the baseline-corrected power spectra of the PSPs from all populations to the excitatory populations.

To further investigate the mechanism behind the locally (fixed power) negative but globally (overall) positive alpha-BOLD correlations observed by (Magri et al., 2012) and reproduced in Figure 6C, we present three example cases in which both global and local alpha-BOLD correlations are negative in the supplementary Figures S4, S5, and S6. A key factor in these cases is the alpha-gamma correlation. All three cases show a negative alpha-gamma correlation, contrasting with the positive alpha-gamma correlation in Figure 6B. A negative alpha-gamma correlation can arise when external inputs travel between clusters (e.g., between clusters 2 and 6), causing alpha-gamma switching (supplementary Figure S4); from reducing common noise in external inputs (Figure 5 vs. supplementary Figure S5); or increasing the alpha-gamma contrast in the spectrogram by increasing the lateral input *I*_lat_ (Figure 5 vs. supplementary Figure S6). These results suggest that the locally negative and globally positive alpha-BOLD correlations found in (Magri et al., 2012) may be associated with a positive alpha-gamma correlation, potentially due to the relatively low *I*_lat_ (reflecting the anaesthetized state of the monkey) or noisy external input.

### Population activities and EEG spectra

We have so far considered the model output in the form of EEG and BOLD. The model, with its detailed structure, also allows us to further investigate the factors that determine the phenomena just reported. We first focus on which neural populations drive the EEG spectra in different frequency bands. In Figure 7, the top panels show the baseline-removed EEG spectra, and the bottom panels show the correlation spectra to elucidate the relationship between neural population activity and EEG power in different frequency bands. Some neural populations (including E6, S6, and VIP across all layers) only have a marginal influence on the EEG spectra. This can be explained by the relatively weak connections from these populations to the E populations, and therefore the PSPs are dominated by synaptic noise (cf. Figure S7C). We note here that these results are obtained in the constant input condition, where the dynamics of the neural populations are driven by independent synaptic noise.

The classical frequency bands relatively well delimit the correlation spectra. The delta and theta bands (prominent in Clusters 5 to 7) highly correlate with E2/3, E5ET, SOM5, and PV populations. In essence, all layers are involved in these slow rhythms. The alpha band (most prominent in Cluster 2) highly correlates with E2/3, E5ET, SOM2/3, SOM5, PV5, and PV6, suggesting that alpha activity is mostly, but not exclusively, driven by neural activity in the infragranular layers. The beta band has a relatively transitional property, for example, with an overlap with theta and alpha bands (in Cluster 2, < 20Hz) and with the gamma band (in Cluster 2, > 20 Hz). The gamma band (strong in Clusters 3, 4, 6, and 7) shows a high correlation with E (2/3, 4, and 5ET) and PV (all layers). SOM2/3 is also involved in a narrow frequency band close to 40Hz, and SOM4 in the extreme gamma case (Cluster 4).

### Population activities and EEG-BOLD correlation

To examine the correspondence between population activities and EEG-BOLD correlation, we calculate the correlations between EEG-BOLD maps (alpha-BOLD and gamma-BOLD correlations, as in Figure 4AB) and population firing rate maps (mean, standard deviation, and narrow-band power; a subset illustrated in supplementary Figure S2CD) across the input space (*I*_lat_ = [2.5–4.5], *I*_mod_ = [0–25], and *I*_tha_ = [0–5]). In Figure 8A, the alpha-BOLD correlation corresponds mainly with the standard deviation of firing rates in a subset of neural populations (e.g., PV4, VIP4, E5, PV5, SOM5, PV6, and SOM6). The delta and theta activities in these populations (as well as E2/3) also show positive correlations with the alpha-BOLD correlation. This highlights a strong correspondence with slow activity in infragranular populations. In Figure 8B, the gamma-BOLD correlation corresponds to both mean and standard deviation of firing rates of neural populations across all layers (e.g., E2/3, PV2/3, SOM2/3, E4, E5ET, PV5, SOM5, VIP5, E6, and VIP6). The gamma activities in these populations also show positive correlations with the gamma-BOLD correlation.

**Figure 8.**
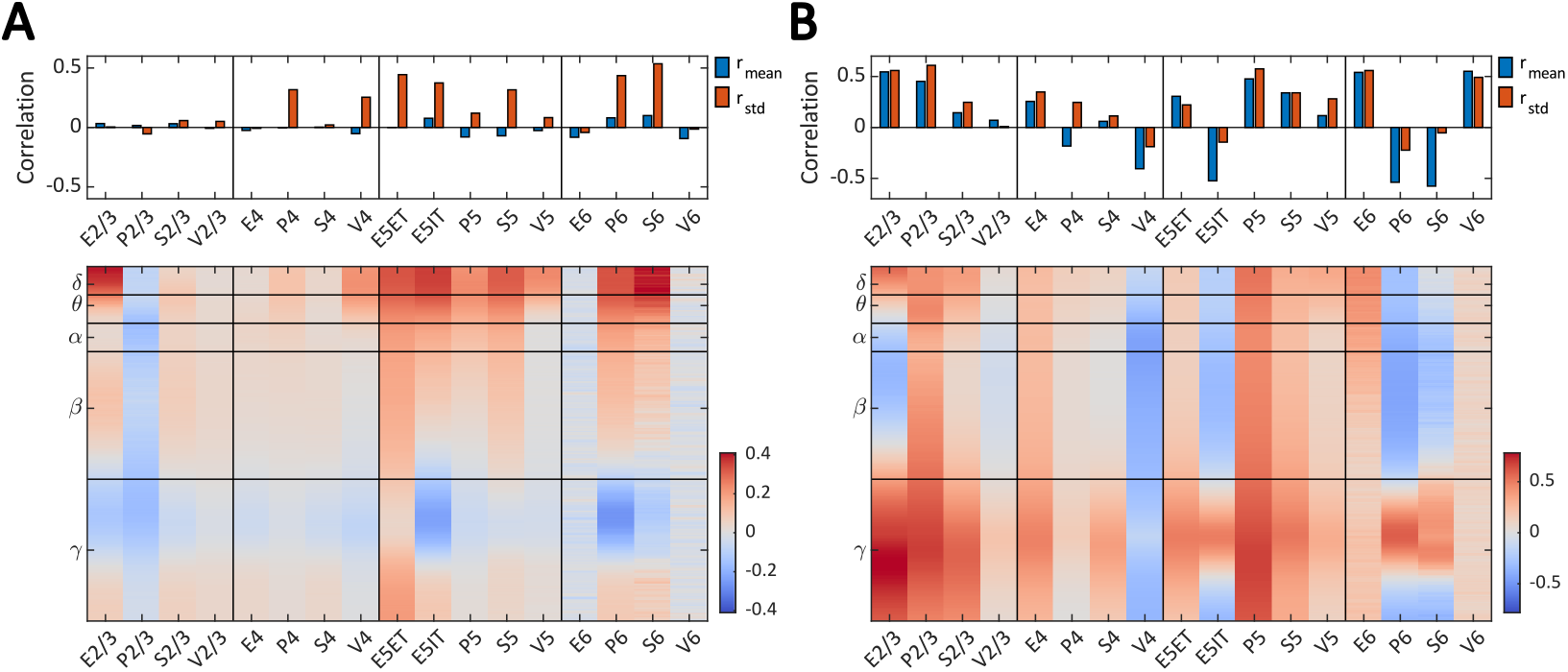
Populational relationship with alpha-BOLD and gamma-BOLD correlations. (**A**) Similarity (measured by correlation across the input space) between alpha-BOLD correlation and various metrics. Top panel: with mean firing rate in blue bars (cf. supplementary Figure S7A) and standard deviation of firing rate in red bars (cf. supplementary Figure S7B) (top row). Bottom panel: with relative power change in firing rate at each frequency bin (cf. supplementary Figure S7B). (**B**) Similarity between gamma-BOLD correlation and the same metrics as in (A).

### Circuit-level effect on BOLD response to optogenetic stimulation

To further validate the model, we aim to reproduce experimental findings on BOLD signal changes evoked by optogenetic stimulation of specific neuronal populations. Optogenetic studies have shown that selective stimulation of inhibitory interneuron sub-classes (PV, SOM, VIP, and nitric oxide synthase [NOS]) can induce distinct changes in cerebral blood flow (CBF) and blood volume, thereby producing positive or negative BOLD responses depending on the cell type and stimulation parameters (Cauli et al., 2004; Krawchuk et al., 2020; L. Lee et al., 2020; Moon et al., 2021; Uhlirova et al., 2016). While such experiments demonstrate that neurovascular factors (e.g., nitric oxide release from NOS interneurons) can directly modulate hemodynamics, we do not explicitly model these vascular mechanisms. Instead, our microcircuit model demonstrates the circuit-level consequences of selectively perturbing inhibitory populations that indirectly shape BOLD responses through changes in excitatory-inhibitory balance, which are unavoidable in optogenetic experiments.

In Figure 9, we selectively activated PV2/3, SOM2/3, or VIP2/3 populations, mimicking optogenetic stimulation conditions in which superficial layers receive stronger light intensity. Simulated transient BOLD responses were examined under a baseline condition [*I*_lat_, *I*_mod_, *I*_tha_] = [4.5, 5, 0], located within the alpha-dominant regime. Activation of PV2/3 led to a decrease in BOLD activity, accompanied by increased firing in PV6 and VIP2/3, and decreased firing in PV5, SOM2/3, and E5ET (Figure 9A). Activation of SOM2/3 also resulted in a negative BOLD response, but with a distinct circuit signature: increased firing in PV6, and reduced firing in PV2/3, PV5, VIP2/3, and E5ET (Figure 9B). In contrast, activation of VIP2/3 produced a positive BOLD response, associated with suppression of SOM2/3 and PV6, and disinhibition-driven increases in PV2/3, PV5, and E5ET firing rates (Figure 9C). For visualization purposes, stimulation intensities were coarsely tuned across conditions to yield comparable BOLD amplitude changes and clearly distinguishable firing-rate patterns.

**Figure 9.**
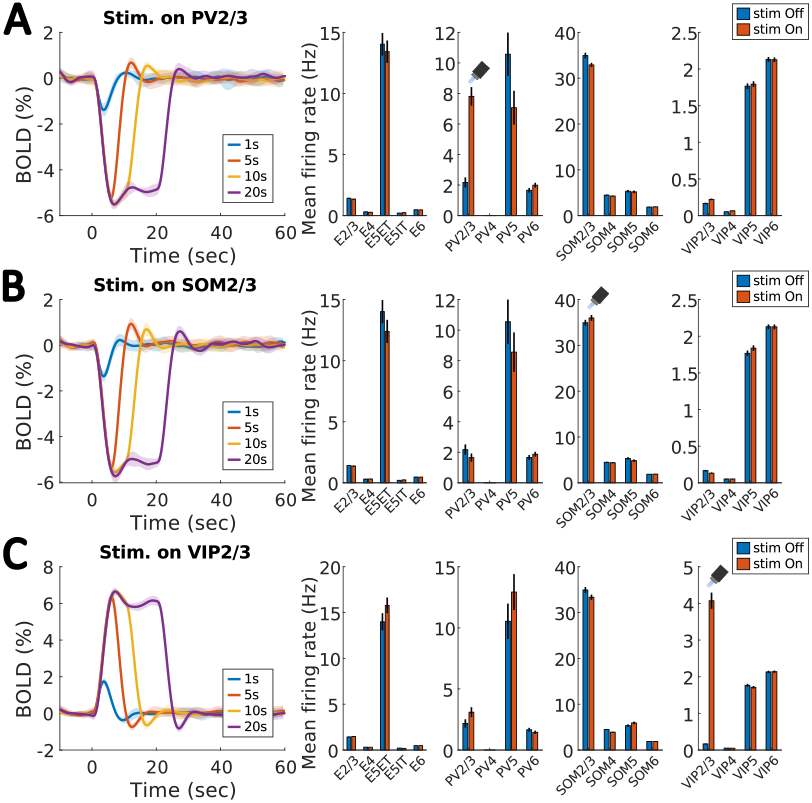
BOLD responses to selective stimulation of interneuron subtypes. Relative changes in simulated BOLD signals are shown for three conditions in which an additional step input (onset = 0 sec; duration = 1, 5, 10, or 20 sec) was applied to one of three interneuron populations: (**A**) PV2/3, (**B**) SOM2/3, or (**C**) VIP2/3. Each condition was simulated 10 times. Colored traces indicate the mean BOLD response, and shaded regions denote the standard deviation across runs. To the right of each BOLD panel, bar plots show the mean firing rates of the 17 neural populations during the stimulation period (“stim. On”) and after stimulation (“stim. Off”), with error bars indicating the standard deviation. Baseline external inputs were set to [*I*_lat_, *I*_mod_, *I*_tha_] = [4.5, 5, 0]. The stimulation intensities were manually tuned (A: *I*_PV2/3_ = 2; B: *I*_SOM2/3_ = 0.3; C: *I*_VIP2/3_ = 7) to produce BOLD changes with comparable amplitudes. The stimulation input itself was not included in the BOLD forward model, mimicking optogenetic stimulation in animal experiments where the optical input does not directly contribute to the hemodynamic signal.

The polarity of the simulated BOLD responses under this exemplary baseline condition broadly aligns with findings reported in the literature. These include SOM interneuron stimulation, which has been associated with negative BOLD responses (Cauli et al., 2004), biphasic BOLD responses (Krawchuk et al., 2020), and negative BOLD signals in surrounding cortical regions (L. Lee et al., 2020), as well as VIP interneuron stimulation, which has been linked to positive BOLD changes (Cauli et al., 2004). In our simulations, however, the polarity of the BOLD response depends strongly on the baseline network state, parameterized by [*I*_lat_, *I*_mod_, *I*_tha_]. Across baseline conditions, stimulation of PV2/3 generally produced negative BOLD responses. In contrast, stimulation of SOM_2/3_ resulted in negative BOLD responses under weak *I*_tha_ but positive BOLD responses under strong *I*_tha_, whereas stimulation of VIP_2/3_ exhibited the opposite dependence (Supplementary Figure S10). These results demonstrate that the same interneuron perturbation can give rise to opposite BOLD polarities depending on the ongoing circuit state, providing a circuit-level explanation for the heterogeneous optogenetic BOLD outcomes observed experimentally. Additional factors, such as stimulation of interneurons in deeper layers (e.g., L4 or L5/6) or geometric effects related to center-surround organization, may further modulate circuit-level responses but are not explored in the present simulations.

## Discussion

In this study, we investigate how long-range inputs to cortical microcircuits contribute to the heterogeneity of EEG-BOLD correlations observed in experimental data (Figure 1A-D). We systematically examine the relationship between EEG rhythms and BOLD amplitudes by scanning the levels of external inputs in a detailed microcircuit model (Figures 2, 3, and S2). We further demonstrate that both constant and varying input conditions can produce positive or negative EEG-BOLD correlations. Under constant input conditions, the model produces weak EEG-BOLD correlations (Figure 4), which may correspond to the observed distribution of alpha-BOLD correlations (Figure 1B). However, time-varying inputs exert a more pronounced effect (Figures S8 and S9), resulting in stronger EEG-BOLD correlations (Figure 5) and better alignment with experimental data (e.g., LFP-BOLD correlation in Figure 6). For a typical case where strong alpha activity and negative alpha-BOLD correlations are observed in occipital regions during resting-state, a time-varying input condition (with moderate *I*_lat_, low *I*_tha_, and low *I*_mod_) offers a more plausible explanation.

### Sources of EEG rhythms

While low- and high-frequency EEG bands are conventionally associated with SOM and PV neurons, respectively (Chen et al., 2017; Onorato et al., 2025; Vierling-Claassen et al., 2010), our findings reveal a more intricate involvement of neural populations in the correlation spectra, where SOM and PV neurons jointly contribute to shifts in the resonance frequency (Figure 7). In addition, we found that low-frequency EEG activity (<20 Hz) was driven more strongly by infragranular than supragranular populations. Conversely, high-frequency EEG activity (>20 Hz) is driven more strongly by supragranular populations. Nevertheless, drivers in both cases emerged from all layers, which doesn’t justify assigning specific EEG frequency bands unequivocally to a specific layer. Furthermore, the same neural population can be involved in both low and high frequency EEG activity, which is particularly the case for pyramidal cells and PV interneurons in layers 2/3 and 5.

### Comparing models of EEG rhythms

Our model generates diverse EEG rhythms by leveraging excitatory-inhibitory feedback loops that resonate with synaptic noise (Hahn et al., 2022; Vierling-Claassen et al., 2010), shaped by variations in firing functions and synaptic time constants across cell types. This mechanism is not in conflict with, but rather complements, other mechanisms proposed for EEG rhythm generation. For example, several studies have explored the roles of different inhibitory interneurons (e.g., PV, SOM, and VIP neurons) in the emergence of oscillations within local microcircuits (Domhof & Tiesinga, 2021; Hahn et al., 2022; B. Lee et al., 2018; Veit et al., 2017). Our model aligns with these works in many respects, such as the disinhibitory role of VIP neurons, the association of fast oscillations with PV-dominated states, and the emergence of slower oscillations in SOM-dominated states. However, these studies predominantly focused on EEG rhythms in higher frequency bands (e.g., beta and gamma, >20 Hz). Other studies, including cortical network models (Nakagawa et al., 2014) and thalamocortical models (Pang & Robinson, 2018), focus on structural connectivity and transmission delays at the brain scale to explain alpha and low-frequency oscillations. While these structural factors may account for inter-subject differences in alpha power, they are less relevant for capturing the dynamic changes in alpha power.

### Comparing models of EEG-BOLD relationship

We identified two computational studies that have attempted to explain alpha-BOLD anticorrelation. In one study, a network model was developed to study EEG-fMRI correlation at the whole-brain level (Schirner et al., 2018). In this study, the source-localised EEG signal is used as a driving input into cortical E-I circuits, and then compared with the model-generated fMRI signal. This study found consistently strong anti-correlations between the alpha power and the fMRI signal, which does not reflect the more nuanced findings of experimental studies. In another study, a thalamocortical model was proposed to address alpha generation and alpha-BOLD anticorrelation (Pang & Robinson, 2018). This thalamocortical model highlights the anticorrelation between alpha power (7.5–13 Hz) and low-frequency power (0–4 Hz), with the latter serving as a proxy for the BOLD signal. This study attributes changes in alpha power to structural factors, such as corticothalamic and intrathalamic feedback strengths. These structural factors are used to explain the alpha-BOLD anticorrelation observed in a group-level analysis. In contrast, our model captures this relationship in the temporal domain, explaining this phenomenon at the single-subject level. Moreover, our model directly simulates the BOLD signal by feeding the sum of EPSPs into the Balloon-Windkessel model, without assuming any low-frequency proxy for BOLD. We conclude that ours is the most realistic model to date that has been used to address the topic of EEG-BOLD correlation.

### Model complexity and limitations

This study examined how external inputs shape the neural dynamics of cortical microcircuits and how these dynamics affect the relationship between two macroscopic primary readouts (i.e., LFP/EEG and BOLD). The cortical microcircuit architecture, together with the EEG and BOLD forward models and the external input configurations, was designed to maintain biological plausibility. Nonetheless, several simplifications were introduced to isolate the key mechanisms of interest, particularly the roles of distinct inhibitory interneuron types and their interactions with excitatory populations that collectively modulate the EEG-BOLD relationship. The implications of these simplifications define the scope and limitations of the present model, as discussed below.

#### (1) Neural dynamics

We considered only AMPA receptor-mediated synaptic transmission (characterized by fast time constants) in excitatory connections, assuming that NMDA receptor activity (with slower time constants) is relatively low during resting states (Jiang et al., 2024). Previous modeling work has shown that realistic local field potential (LFP) spectra can be reproduced using only fast AMPA synapses in reduced cortical microcircuit models (Sanchez-Todo et al., 2023), whereas NMDA receptor dynamics become more prominent during slower, task-related neural processes (Jiang et al., 2024). Similarly, short-term synaptic plasticity (STP) was not included in this study, as previous findings indicate that ongoing spontaneous activity remains qualitatively similar with or without STP (Jiang et al., 2024). Consequently, the present simulation results, including the spectral characteristics of the microcircuits and the modeled EEG-BOLD relationships, should be interpreted as representative of resting-state conditions. To extend this framework to evoked responses or cognitive task states, both NMDA receptor-mediated transmission and STP would need to be incorporated, and their inclusion would likely necessitate a re-evaluation of the resulting spectral properties and EEG-BOLD correlations.

#### (2) LFP/EEG forward model

The LFP/EEG forward model was implemented as a weighted sum of excitatory and inhibitory synaptic activities across neural populations, with the weights representing the amplitude and orientation of current dipoles. Neuronal morphology and the precise spatial distribution of synaptic contacts, typically modeled in biophysically detailed simulations, were not considered here. This simplification should not qualitatively alter the frequency profile of LFP/EEG power or the EEG-BOLD correlation, a conclusion partially supported by tests using randomized dipole orientations (Figure S3). For a more mechanistic understanding, future work could integrate cell-type- and depth-specific information into the forward model. Incorporating the spatial distribution of synaptic locations and depth-dependent dipole positions would allow more precise quantification of how individual cell types contribute to the observed LFP/EEG spectra and their relationship to BOLD (Næss et al., 2021).

#### (3). BOLD forward model

We simulated the BOLD signal using a conventional Balloon-Windkessel model (Friston et al., 2000), which considers only excitatory synaptic activity as its input. Direct contributions of inhibitory neurons to hemodynamic responses were not included. However, emerging evidence suggests that inhibitory interneurons can directly influence cerebral hemodynamics. For instance, activation of nitric oxide synthase (nNOS)-expressing interneurons leads to the production of nitric oxide (NO), which rapidly diffuses to nearby vascular smooth muscle cells and pericytes, causing vasodilation and increased BOLD signal (Cauli et al., 2004; Krawchuk et al., 2020; L. Lee et al., 2020; Moon et al., 2021; Vazquez et al., 2018). We assume that incorporating nNOS interneurons would not qualitatively change the sign of the simulated EEG-BOLD correlations, since these cells receive broad excitatory inputs from pyramidal neurons and thus primarily reflect local excitatory activity– the same activity currently driving the Balloon-Windkessel model. Other inhibitory populations, such as SOM and VIP interneurons, are also known to modulate BOLD responses (Cauli et al., 2004; Krawchuk et al., 2020; L. Lee et al., 2020; Uhlirova et al., 2016). Their indirect vascular influences depend on how they shape downstream circuits, for example, SOM-mediated surround inhibition and VIP-mediated disinhibition. We illustrate these circuit-level effects in Figure 9, where selectively stimulating interneuron subtypes leads to a reorganization of population activity and results in either positive or negative BOLD changes. These effects are implicitly captured through the microcircuit input configuration used in our simulations. However, the direct neurovascular effects of these interneurons were not explicitly modeled. Many additional mechanisms of neurovascular and neurometabolic coupling influence local BOLD signals (see (Howarth et al., 2021; Jolivet et al., 2015)), but these lie beyond the current scope. Future work could compare the present framework against fine-tuned BOLD forward models to quantify the relative impact of such mechanisms on EEG-BOLD coupling.

#### (4). Input configuration

External inputs were simplified into three distinct, non-overlapping, and temporally independent types. This configuration was designed to provide mechanistic insight into model behavior rather than to reproduce detailed resting-state statistics. Our results demonstrate that varying external input parameters can account for heterogeneity in EEG-BOLD correlations (Figure 1). Future large-scale (whole-brain) models could incorporate additional biological sources of variability, such as regional differences in PV/SOM interneuron density or cortical hierarchical level, to more accurately fit individual EEG-BOLD data.

Despite these simplifications, the current approach offers a tractable yet biologically grounded framework for understanding how neural population dynamics give rise to EEG-BOLD correlations. Subsequent model extensions that incorporate the factors discussed above would broaden its applicability and enhance its predictive accuracy.

### Model predictions

The microcircuit model makes tangible predictions, and its scalability makes it an ideal candidate for further extensions. A simple prediction is concerned with the variability of the EEG-BOLD correlation when it is computed from finite time series. The model provides evidence that this variability can be decreased by increasing the length of the time series (a four-fold increase halves the standard deviation). While longer recordings would be more costly in their acquisition, they would allow the detection of more subtle correlations that are otherwise missed. Longer recordings would also enable the test of another prediction the model makes: the power in the delta and theta band (Figure 2A) correlates with the standard deviation of the BOLD signal (Figure S2C).

Our investigation focused on different forms of external input that target distinct neural populations. The results could be tested experimentally in a mouse model using optogenetics to target specific neural populations, such as VIP interneurons to mimic modulatory input. The model in turn could also be used to help design optimal stimulation protocols for a variety of applications, including the effect of specific neural populations on LFP and BOLD.

## Methods

### Cortical microcircuit model

The cortical microcircuit model comprises 17 neural populations, including excitatory (E) neurons and three types of inhibitory interneurons—parvalbumin-positive (PV), somatostatin-positive (SOM), and vasoactive intestinal peptide-positive (VIP) interneurons—distributed across cortical layers 2/3, 4, 5, and 6 (Figure 1E). The connectivity *w*_*ji*_ from population *i* to population *j* is derived from the connection probability *p*_*ji*_, synaptic strength *s*_*ji*_, and cell count *c*_*i*_ of source population *i* (Campagnola et al., 2022; Fang et al., 2022; Gonchar et al., 2008). The cell count *c*_*i*_ is normalised to total cell counts so thatn 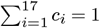. The final connectivity matrix *W*, consisting of these pairwise connectivity values *w*_*ji*_, is rescaled by a global scaling factor *G* (corresponding to the number of neurons to be considered in the cortical microcircuit model) to ensure proper model behavior. In this study, we use *G* = 20, 000, where the model fluctuates with the synaptic noise without entering oscillatory states in the defined external input range.

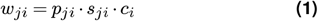

The dynamics of the cortical microcircuit model are described by two variables: firing rate *m*(*t*) and PSP *v*(*t*), and the evolution is governed by potential-to-rate and rate-to-potential operators (Chien et al., 2023; Jansen & Rit, 1995; Spiegler et al., 2010; Wang & Knösche, 2013). The potential-to-rate operator translates the PSP *v*(*t*) of a population into its firing rate *m*(*t*) using a sigmoid function:

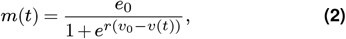

where *e*_0_ is the maximum firing rate (Hz), *v*_0_ is the potential threshold (mV), and *r* determines the steepness of the sigmoid curve. These parameters (listed in Table 1) are derived by fitting the sigmoid function to the mean F-I curve from the database of single cells from Mouse Brain of Allen Brain Atlas (https://celltypes.brain-map.org/) for each neuron type (E, PV, SOM, and VIP).

**Table 1.** Parameter setting for potential-to-rate operators.

| | $e_0$ (Hz) | $v_0$ (mV) | $r$ (1/mV) |
| --- | --- | --- | --- |
| E | 31.40 | 32.11 | 0.13 |
| PV | 166.83 | 40.03 | 0.14 |
| SOM | 56.95 | 42.02 | 0.08 |
| VIP | 38.53 | 37.86 | 0.07 |

The rate-to-potential operator characterises the transmission of firing activity *m*_*i*_(*t*) from a source population *i* to the PSP *v*_*ji*_(*t*) of a target population *j*. This process accounts for dendritic and synaptic properties and is described by a second-order ordinary differential equation:

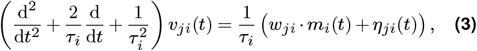

where *τ*_*i*_ is the time constant that depends on the source population *i, w*_*ji*_ is the connectivity, and *η*_*ji*_(*t*) is the synaptic noise. In this study, we use *τ*_ext_ = 6 msec, *τ*_E_ = 6 msec, *τ*_PV_ = 3 msec, *τ*_SOM_ = 20 msec, and *τ*_VIP_ = 15 msec for the connections from external input and source populations E, PV, SOM, and VIP, respectively.

The synaptic noise *η*(*t*), zero-mean white Gaussian noise with standard deviation *σ*_*η*_, is independently applied to all internal and external input connections. In this study, we use *σ*_*η*_ = 1 mV. Lower noise levels (e.g., *σ*_*η*_ < 1 mV) result in similar diverse spectral properties with a shift in the power spectrum, whereas higher noise levels (e.g., *σ*_*η*_ ≥10 mV) lead to reduced signal-to-noise ratios (the signal being the PSPs from neural populations and external inputs) and diminish the diversity.

### External inputs

The cortical microcircuit model is driven by three distinct types of external inputs: lateral input *I*_lat_, modulatory input *I*_mod_, and thalamic input *I*_tha_, forming a three-dimensional input space for exploring model dynamics (Figure 1E). Each input targets specific neural populations within the cortical microcircuit, reflecting simplified interactions observed in real-world neural systems. The lateral input *I*_lat_ is directed to E2/3 and SOM2/3 populations, representing interactions with neighboring cortical microcircuits (Yavorska & Wehr, 2016). The modulatory input *I*_mod_ is delivered to VIP2/3 populations, simulating top-down signals originating from higher-order brain areas (Bastos et al., 2023). Meanwhile, the thalamic input *I*_tha_ targets E4 and PV4 populations, a reduced abstraction of thalamic projections, which in reality also reach L2/3, L5 and L6 populations in a more detailed manner (Ji et al., 2016). This specific input configuration balances simplicity and biological accuracy.

### Forward models

The forward models take PSPs to simulate EEG and BOLD signals (Figure 1F). These simulations are based on distinct biophysical principles that link neural activity to macroscopic brain signals.

The simulation of EEG signals relies on the understanding that EEG signals primarily originate from transmembrane currents at the aligned long dendrites of pyramidal cells (Biasiucci et al., 2019). This corresponds to the excitatory postsynaptic potentials (EPSPs) and inhibitory postsynaptic potentials (IPSPs) within the excitatory populations (E2/3, E4, E5IT, E5ET, and E6). Parameters such as dipole configuration *d*_*ji*_ and cell count *c*_*j*_ of the target excitatory population *j* are incorporated to weigh the contribution of these currents to the EEG signal.

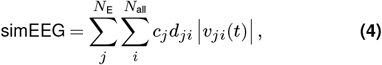

where *v*_*ji*_(*t*) corresponds to the transmembrane current at the target E population *j* introduced by source population *i, d*_*ji*_ controls the length and direction of the current dipole, and *c*_*j*_ denotes the cell count of the target E population *j*. In this model, *N*_*E*_ includes 5 excitatory populations, and *N*_*all*_ includes 17 populations and the external input. We set dipole configuration [*d*_E_→E, *d*_PV_ →_E_, *d*_SOM_→ _E_, *d*_VIP_ → _E_, *d*_ext_ → _E_] = [−1, −0.1, 1, 1, 0.1] to roughly catch the distinct distribution of synapses on the pyramidal cells from different types of presynaptic populations. For example, PV targets the cell body of E, and SOM targets the dendrites of E. A more accurate estimation of *d* would rely on the simulation of a cortical microcircuit model consisting of neurons with detailed morphology, such as (Næss et al., 2021).

The simulation of the BOLD signal is based on the assumption that energy consumption is primarily driven by the synaptic activity of excitatory neurons (Howarth et al., 2012). The neural activity signal *z* is therefore calculated as the sum of EPSPs *v*(*t*) of all populations (weighted by cell counts *c* in Table 2) introduced by the excitatory populations and external inputs. The signal *z* is then fed to the Balloon-Windkessel model (Friston et al., 2000, 2003), which captures the complex vascular and metabolic processes underlying the generation of the BOLD signal.

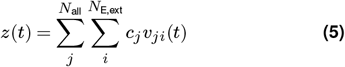

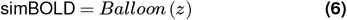

**Table 2.**
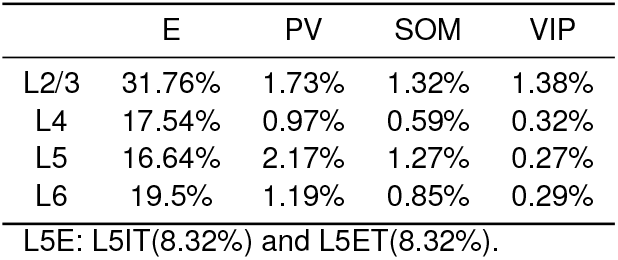
Cell count of populations (in %).

|  | E | PV | SOM | VIP |
| --- | --- | --- | --- | --- |
| L2/3 | 31.76% | 1.73% | 1.32% | 1.38% |
| L4 | 17.54% | 0.97% | 0.59% | 0.32% |
| L5 | 16.64% | 2.17% | 1.27% | 0.27% |
| L6 | 19.5% | 1.19% | 0.85% | 0.29% |

L5E: L5IT(8.32%) and L5ET(8.32%).

### Simulations under constant input condition

For simulations under the constant input condition, the average alpha-BOLD and gamma-BOLD correlations were calculated over 100 runs of 10-minute simulations for each combination of external inputs. The input space was explored with a step size of 1 within the ranges *I*_lat_=[2.5–4.5], *I*_mod_=[0–25], and *I*_tha_=[0–5]. In this input range, the model maintained a fixed-point state in the absence of synaptic noise *η*.

The preprocessing steps for EEG-BOLD correlation analysis adhered to the procedures commonly used in empirical studies. The alpha (8-12 Hz) and gamma (30-45 Hz) envelopes of the simulated EEG were extracted and input into the Balloon-Windkessel model to generate EEG-derived signals. Bandpass filtering (0.008-0.09 Hz) was then applied to both the EEG-derived signals and the simulated BOLD signal. Finally, Pearson correlations were calculated during the stable interval (200-550 seconds) for each simulation run.

### Simulations under varying input condition

For simulations under the varying input conditions, correlations between BOLD signals and the power of each frequency bin (0.5–100 Hz) were calculated across 100 runs of 10-minute simulations. In this setup, the external inputs *I*_lat_, *I*_mod_, and *I*_tha_ were random-walk time series independently generated for each simulation run. The random-walk time series were pre-saved and rescaled to a defined range within the input space for reproducibility.

Preprocessing of these simulations adhered to methods described by (Magri et al., 2012). The time series of overall LFP power was defined as the sum of the simEEG spectrogram over the 0.5–100 Hz frequency range. A fixed-LFP power time series was extracted at defined percentile intervals (0–100%, in steps of 10%) of the overall LFP power. When correlating these time series with simBOLD, a delay of 3.5 seconds was introduced to account for hemodynamic response lag.

## Code availability

All code used in this study is openly available on GitHub at the following link: https://github.com/vscChien/EEGBOLDmodeling.

## Acknowledgments

This work was supported by a Lumina-Quaeruntur fellowship (LQ100302301 awarded to H.S.) founded by the Czech Academy of Sciences, the Czech Science Foundation project (No. 21-32608S), and the Brain dynamics project (No. CZ.02.01.01/00/22_008/0004643) funded by the European Regional Development Fund. We acknowledge the core facility MAFIL supported by the Czech-BioImaging large RI project (LM2018129 funded by the Ministry of Education, Youth and Sports of the Czech Republic) for their support with obtaining scientific data presented in this paper. Computational resources were provided by the e-INFRA CZ project (ID:90254), supported by the Ministry of Education, Youth and Sports of the Czech Republic.

## Author contributions

Conceptualization: V.C., S.J., T.K., J.H., and H.S.; Data curation & formal analysis: S.J.; Methodology & software: V.C.; Funding acquisition: H.S. and J.H.; Supervision: H.S., T.K., and J.H.; Writing - original draft: V.C., S.J., and H.S.; Writing - review & editing: V.C., S.J., T.K., J.H., and H.S.

## Competing interests

The authors declare no competing interests.

## Supplementary Methods

### Empirical EEG-fMRI dataset

For the qualitative comparison of the proposed model behavior in terms of alpha-BOLD correlation, we analyzed a large 72 subjects (mean age: 31.4, range: 18.3 – 50.6, 36 males and 36 females) empirical EEG-fMRI data set acquired in the Central European Institute of Technology (CEITEC) in Brno, Czech Republic.

### Data acquisition and preprocessing

Participants underwent a 20 minutes eyes-closed resting-state EEG-fMRI recording session. Specific details about the data acquisition and preprocessing can be found in our previous study (Jiricek et al., 2025). In short, the functional MRI was acquired via a multiband multi-echo EPI sequence (TR = 650 ms, TE = 14.60/33.56/52.52 ms, 48 slices, slice thickness 3 mm). EEG data were recorded at 1000 Hz sampling frequency using a 256-channel MR-compatible EGI Hydrocell electrode net, along with ECG and breathing signals. Functional MRI data were preprocessed using SPM12 (Penny et al., 2011) and in-house scripts, including spatial realignment, echo fusion via tSNR-weighted averaging, and artifact regression with RETROICOR (Glover et al., 2000) using ECG and breathing signals. Additional preprocessing included orthogonalizing the grey matter signal to potential artifacts (e.g., head motion, white matter/CSF PCA components) and applying a low-pass filter (cut-off 0.09 Hz). The voxel-wise grey matter BOLD time series, originally at a voxel size of 3×3×3 mm, were interpolated using nearest-neighbor interpolation to align with the 6mm regular grid of the EEG source model (described further in the text). This step was necessary to be able to compute local EEG alpha-BOLD coupling. Raw EEG data were preprocessed using a fully-automated pipeline (Liu et al., 2017, 2018; Marino et al., 2019) with MATLAB functions and toolboxes such as SPM (Penny et al., 2011), Fieldtrip (Oostenveld et al., 2011), and EEGLAB (Delorme & Makeig, 2004) toolboxes, starting with main artifacts removal (FASTR for gradient artifacts (Niazy et al., 2005) and adaptive optimal basis set (Marino et al., 2018) for ballistocardiogram artifacts). Bad channels were identified based on low correlation and high variance in non-physiological frequency bands and interpolated using neighboring channels. EEG data were then filtered (1–80 Hz), and ICA (FastICA) was applied to remove artifacts like EOG and EMG based on correlation with reference signals, power spectrum similarity to 1/f, and kurtosis. Finally, we filtered the EEG data to alpha frequency band (8–12 Hz) and rereferenced to average reference.

### Alpha-BOLD correlation

Since we model local microcircuit dynamics, we decided to compute alpha-BOLD correlation on the source-level EEG signals obtained by the very same analytical pipeline as in (Jiricek et al., 2025). Briefly, to estimate brain activity sources, we implemented a source localization pipeline using individual T1 images segmented into 12 compartments (Taberna et al., 2021), followed by head model creation. The source model was generated in the brain grey matter with a grid size of 6 mm. The leadfield matrix was computed using a finite element method (FEM) solver, and sources were estimated with the eLORETA inverse algorithm (Pascual-Marqui et al., 2011). To estimate a single EEG envelope time series, we combined dipole moment components in the three Cartesian axes by assuming freely rotating dipole directions as in (Jiricek et al., 2025). We further followed typical processing steps, including downsampling the EEG envelope to the BOLD TR, convolution with the canonical HRF, and bandpass filtering (0.008–0.09 Hz). For each subject and source model position, the BOLD time series was correlated with the alpha regressor using Pearson’s correlation. All visualizations were performed using the BrainNet Viewer tool (Xia et al., 2013).

## Supplementary Figures

**Figure S1.**
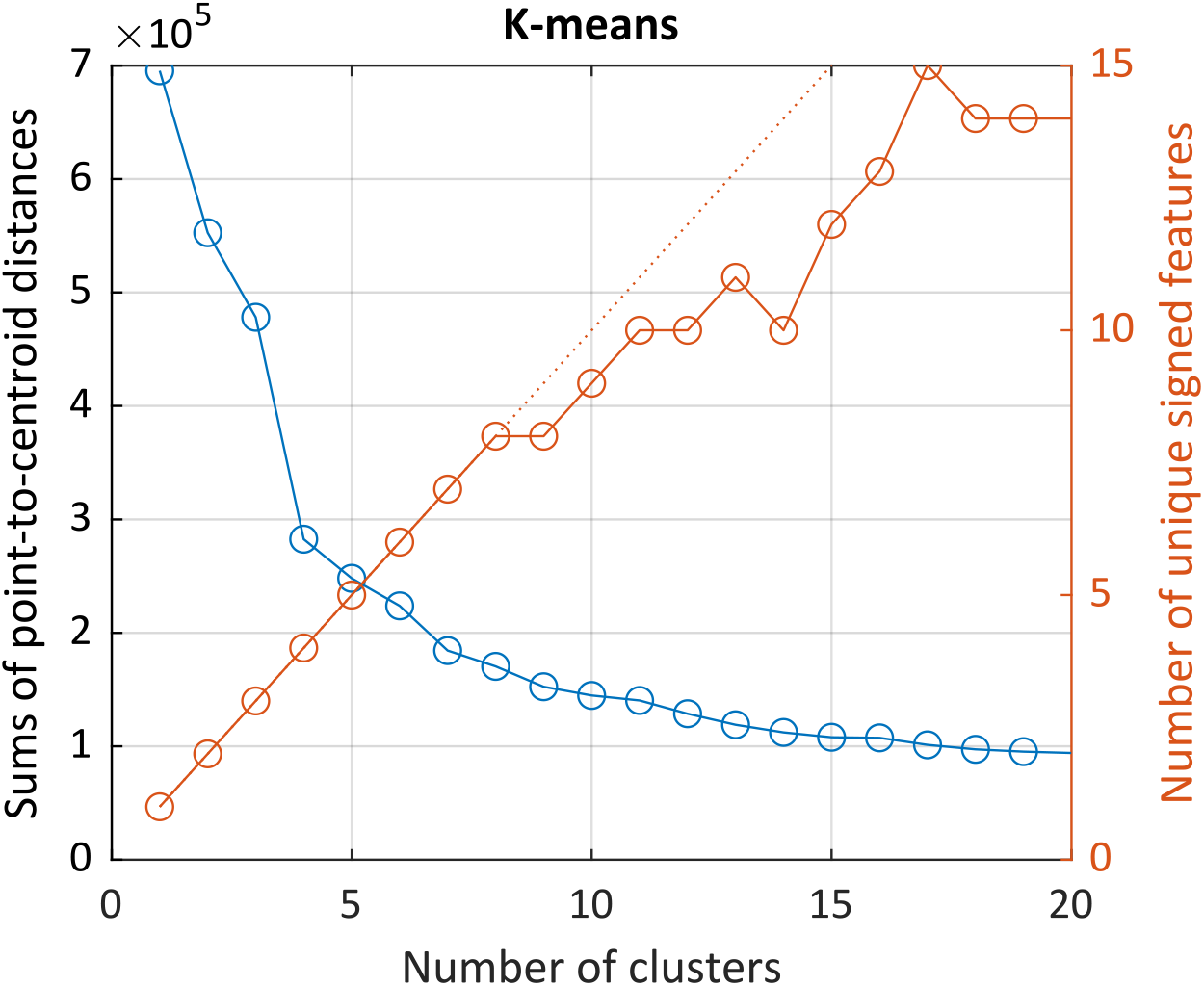
Determining the optimal number of K-means clusters. The K-means algorithm is applied to cluster z-scored power spectra (0.5–50 Hz) within the input space. The blue curve indicates the sum of point-to-centroid distances as a function of the number of clusters, *k*. The red curve represents the number of unique signed features across different values of *k*. A signed feature is defined as a [5*×*1] boolean vector that reflects the direction of mean power change (0: decrease, 1: increase) across delta, theta, alpha, beta, and gamma frequency bands, relative to the baseline. Choosing *k* = 8 maximizes the number of clusters without duplicating signed features while keeping the sum of point-to-centroid distances low.

**Figure S2.**
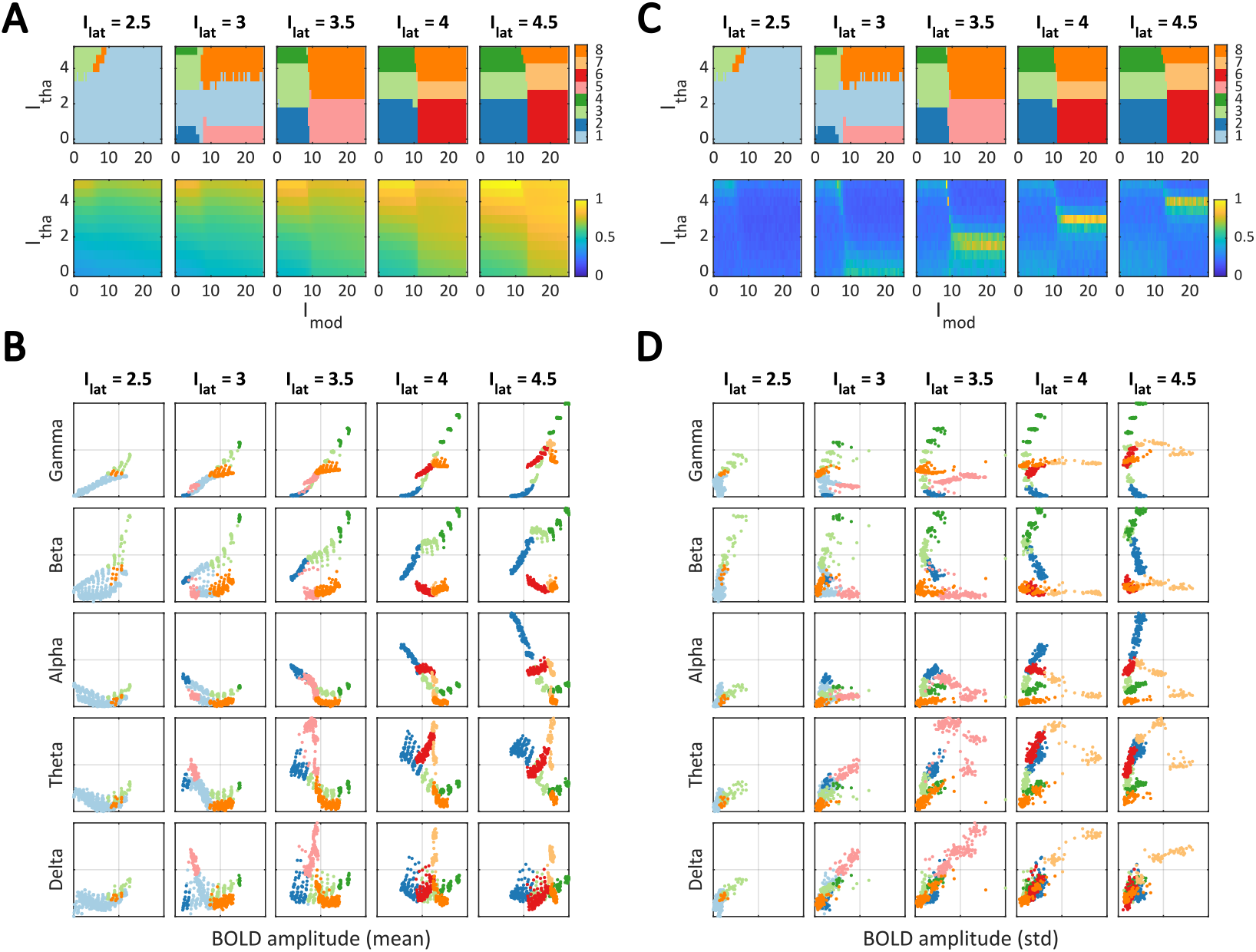
Relationship between EEG bands and BOLD (mean and standard deviation). (**A**) K-means clusters (top row, same as in Figure 2B) and the mean BOLD amplitude (bottom row). (**B**) Scatter plots showing the relationship between gamma, beta, alpha, theta, and delta EEG band powers (from top to bottom) and the mean BOLD amplitude. Both mean BOLD amplitude (x-axis) and EEG band powers (y-axis) are normalized to the range [0–1]. The scatter plots and clusters are represented using the same color scheme. (**C, D**) Same as (**A**) and (**B**), but for the standard deviation (std) of the BOLD amplitude instead of the mean.

**Figure S3.**
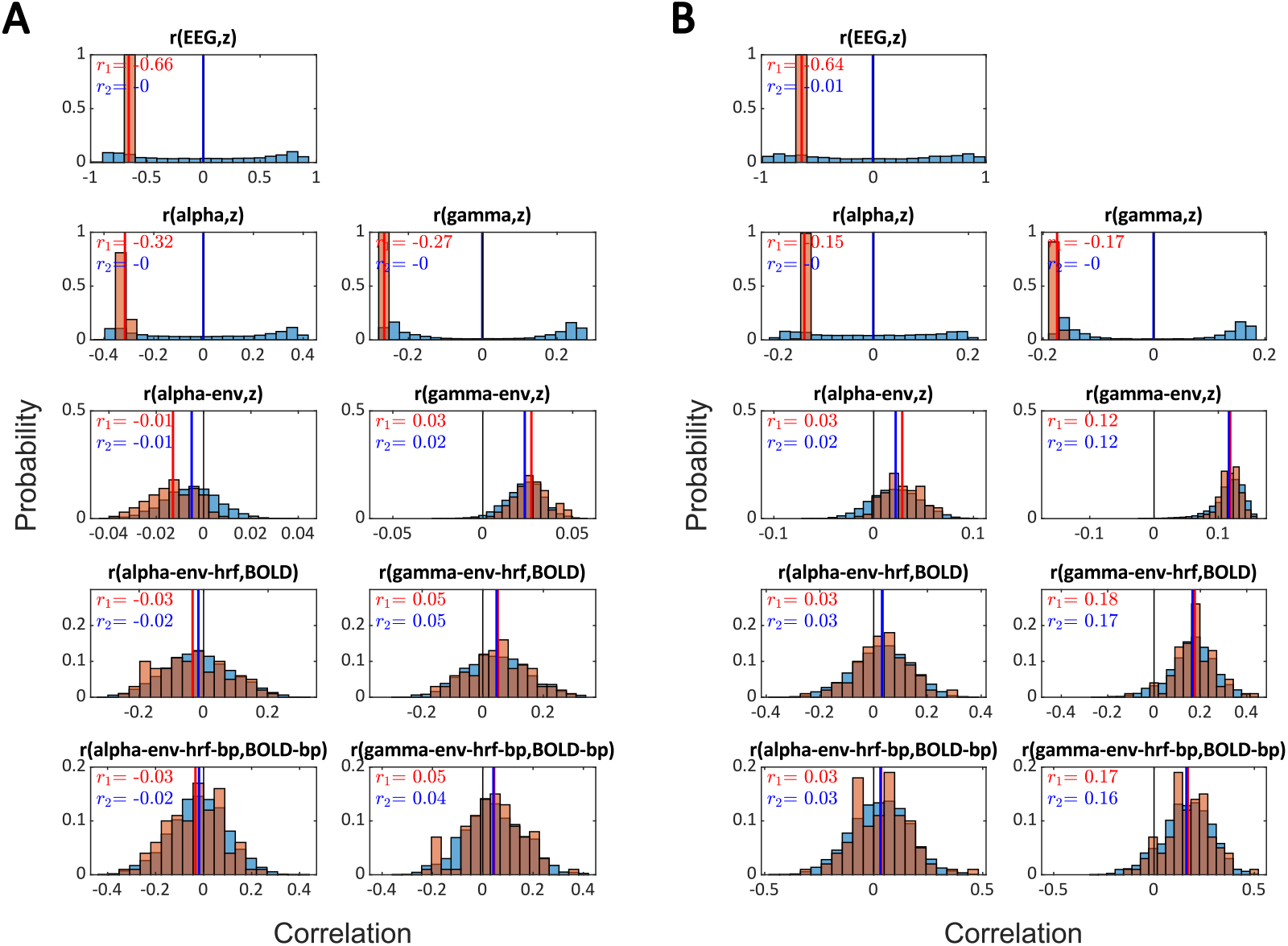
Analysis of dipole effect on EEG-BOLD correlation under constant input conditions. This analysis examines the two input conditions from Figure 4B: (**A**) input condition [*I*_lat_, *I*_mod_, *I*_tha_] = [4.5, 17, 0.5], and (**B**) input condition [*I*_lat_, *I*_mod_, *I*_tha_] = [4.5, 14, 4]. The correlation distribution *r* is calculated across 100 simulations of 10 minutes, each using the default dipole setting (orange histogram, with mean correlation *r*_1_ shown by the red vertical line) and 100 randomized dipole settings per simulation (totaling 10,000 simulations, blue histogram, with mean correlation *r*_2_ shown by the blue vertical line). Abbreviations: ‘EEG’ denotes the simulated EEG signal, ‘z’ refers to the neuronal input for the Balloon-Windkessel model, ‘alpha’ and ‘gamma’ represent the band-pass filtered signals at 8–12 Hz and 35–45 Hz, respectively, ‘-env’ indicates the envelope (amplitude) of the filtered signal, ‘-hrf’ is the hemodynamic response function output of the Balloon-Windkessel model, and ‘-bp’ is the band-pass filtered signal at 0.008–0.09 Hz.

**Figure S4.**
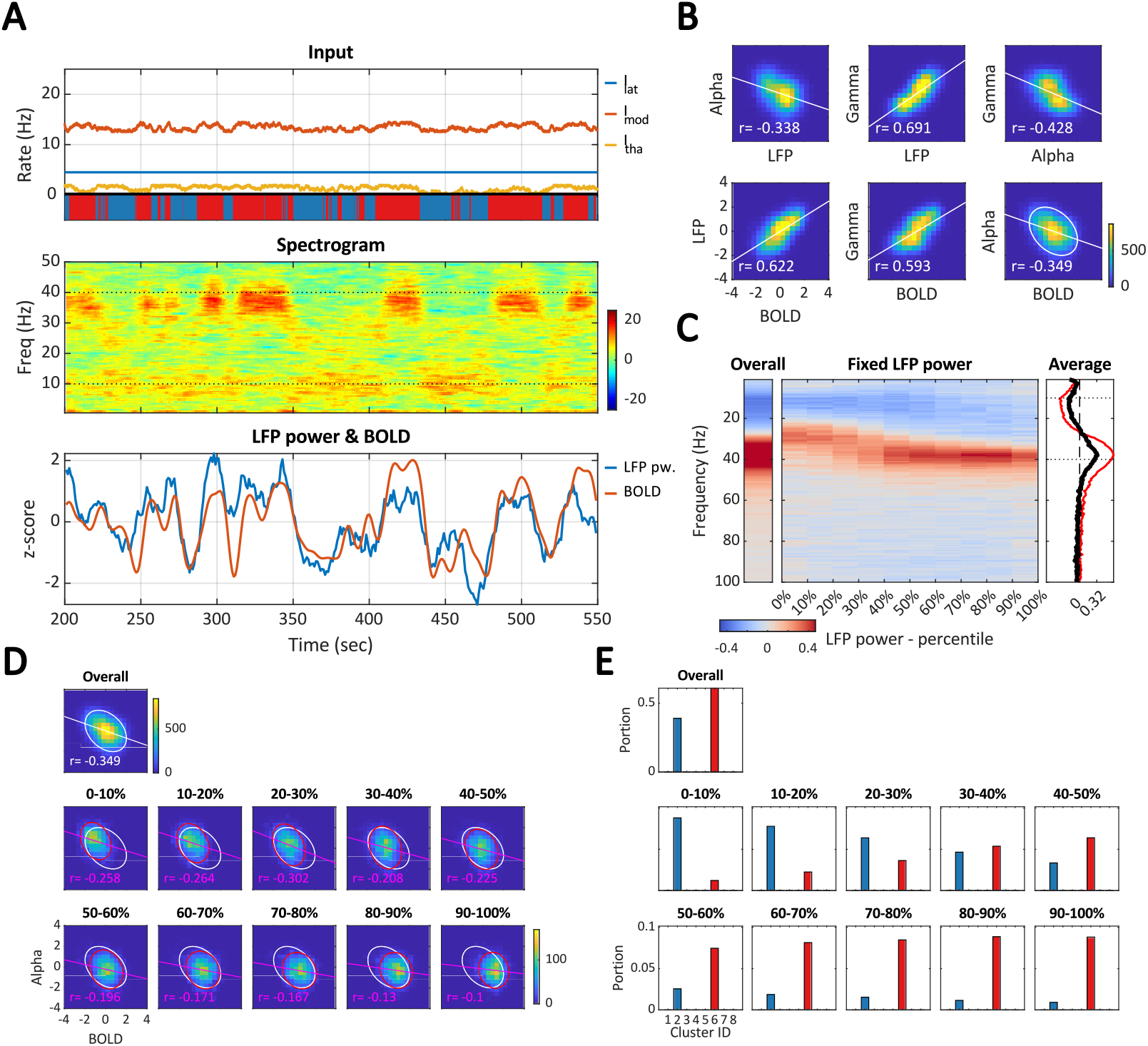
Example 1: alpha-gamma switching. This example uses varying inputs confined to the same range as in Case 2 in Figure 5A: *I*_lat_ = 4.5, *I*_mod_ = [12.5–14.5], and *I*_tha_ = [0–2]. (**A**) The inputs primarily alternate between Clusters 2 (blue stripes) and 6 (red stripes), leading to alpha-gamma band switching in the spectrogram. (**B**) The alpha-LFP correlation, gamma-alpha correlation, and alpha-BOLD correlation are all negative (cf. Figure 6B). (**C**) The global (overall, red curve) and the average local (fixed-power, black curve) correlations are aligned. (**D**) Both global and local correlations are consistently negative. (**E**) Cluster 6 contributes more to higher LFP power than Cluster 2.

**Figure S5.**
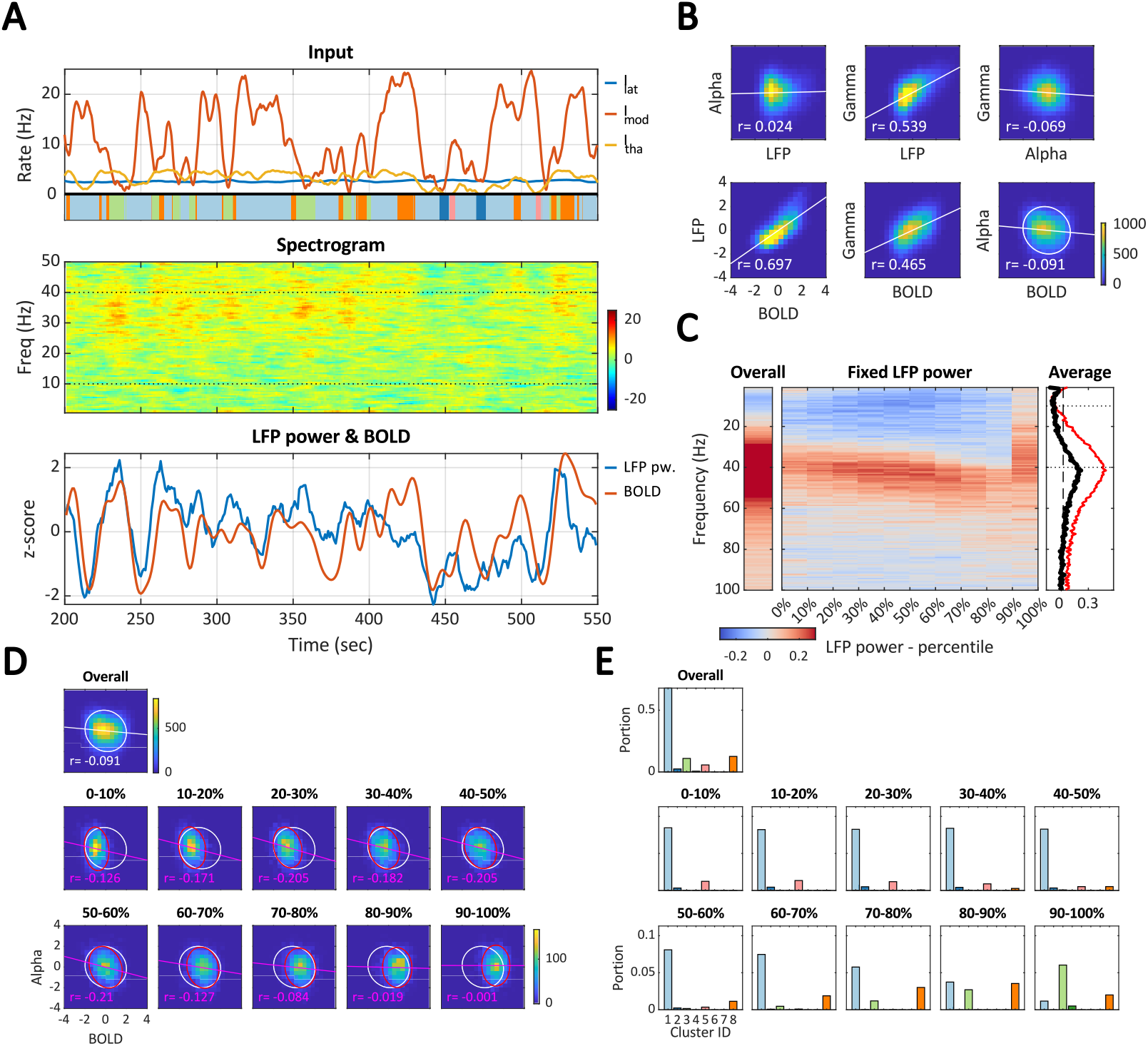
Example 2: smooth external input. This example uses smoothed, varying inputs within the same range *I*_lat_ = [2.5–3], *I*_mod_ = [0–25], and *I*_tha_ = [0–5] as in Figure 6. (**A**) and (**E**) visually resemble Figure 6A and E. (**B**) However, in this case, the gamma-alpha correlation and alpha-BOLD correlation are negative (cf. Figure 6B). (**C**) As a result, both the global (overall, red curve) and mean local (fixed-power, black curve) correlations in the alpha band are negative (cf. Figure 6C). (**D**) Both global and local correlations are consistently negative.

**Figure S6.**
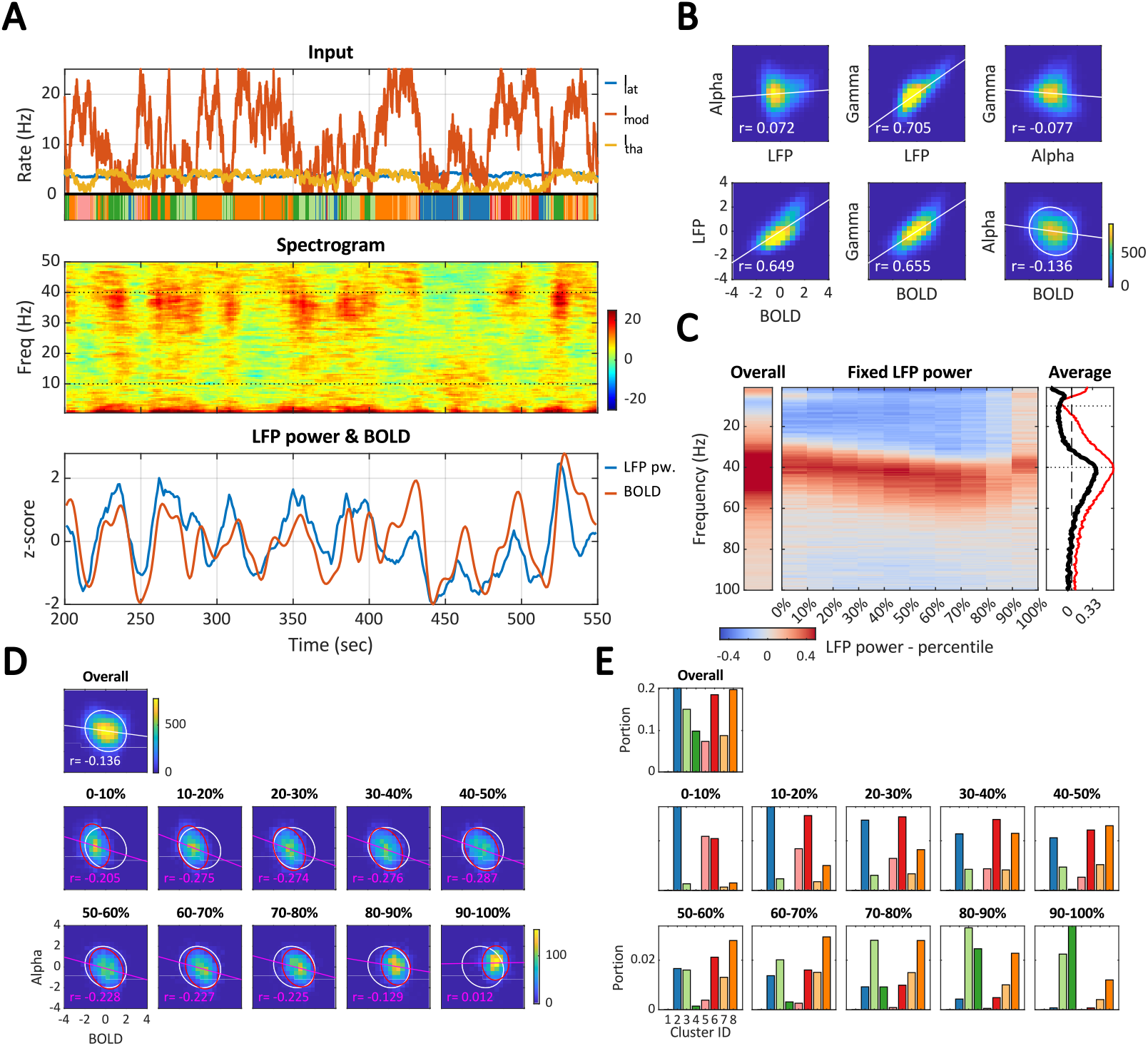
Example 3: increased lateral input. This example uses stronger *I*_lat_ compared to Figure 6: *I*_lat_ = [3.5–4.5], *I*_mod_ = [0–25], and *I*_tha_ = [0–5]. (**A**) and (**E**) The inputs cover a greater number of clusters compared to Figure 6A and E, and the spectrogram displays stronger contrast between spectral patterns due to the increased lateral input. (**B**) In this scenario, the gamma-alpha correlation and alpha-BOLD correlation are negative (cf. Figure 6B). (**C**) Consequently, both the global (overall, red curve) and mean local (fixed-power, black curve) correlations in the alpha band are negative (cf. Figure 6C). (**D**) Both global and local correlations are consistently negative.

**Figure S7.**
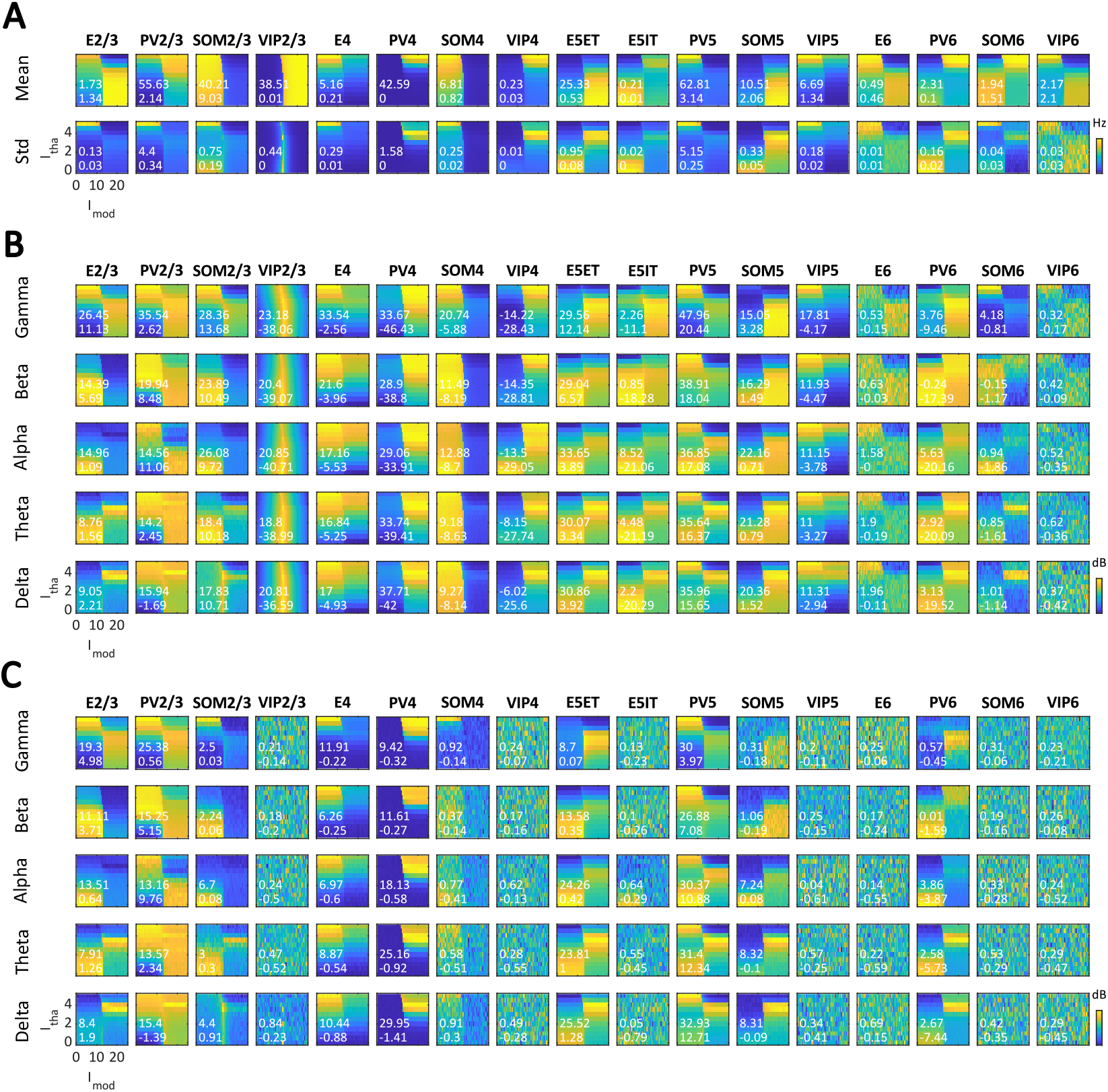
Population-level firing rates and power spectra. (**A**) Mean (top row) and standard deviation (bottom row) of firing rates at input range *I*_lat_ = 4.5, *I*_mod_ = [0–25], and *I*_tha_ = [0–5]. (**B**) Baseline-corrected power at five frequency bands (delta, theta, alpha, beta, and gamma, arranged from bottom to top rows) of the firing rates. (**B**) Baseline-corrected powers of the PSPs from all populations to the excitatory populations. The white text indicates max and min values.

**Figure S8.**
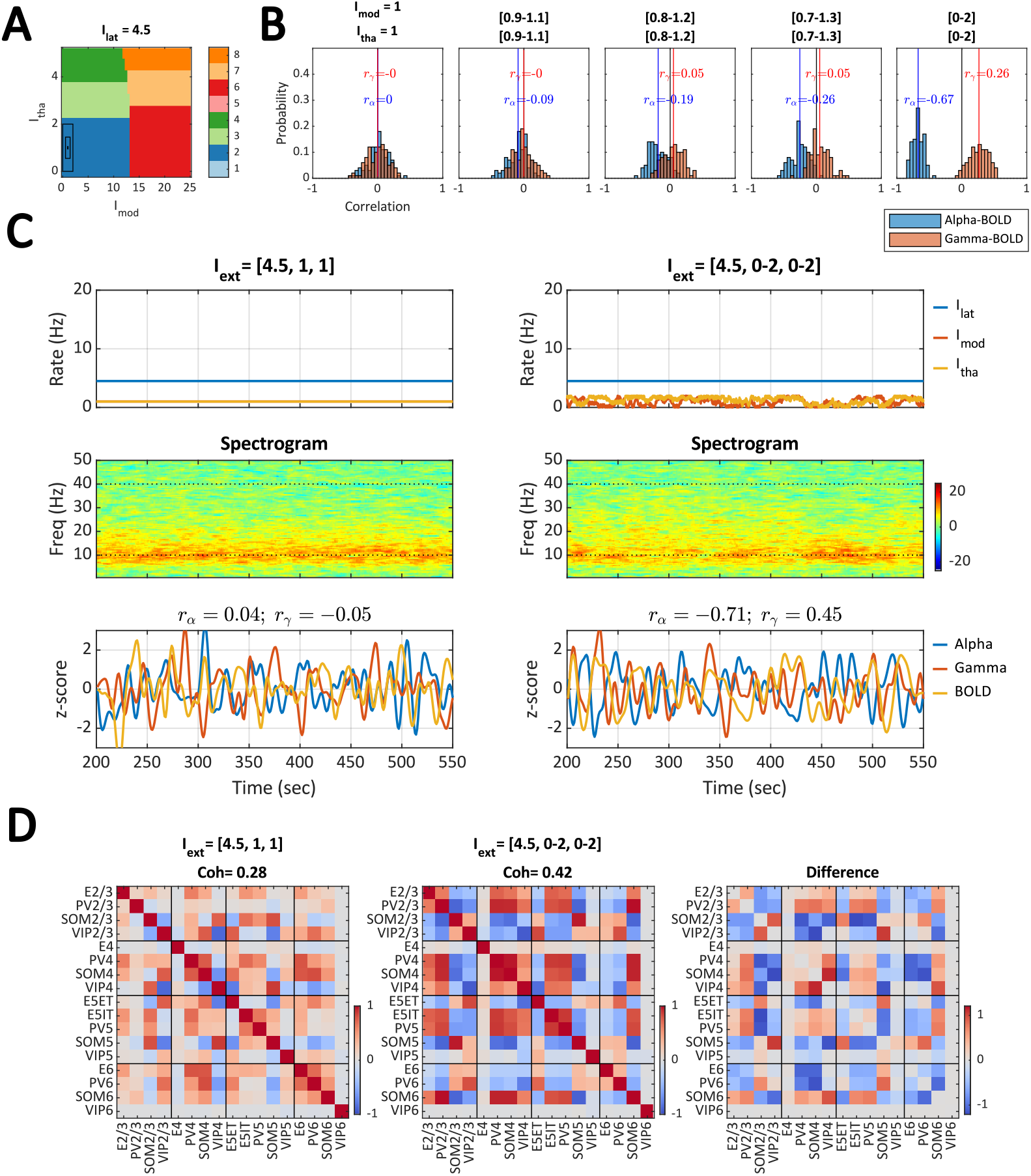
Effect of external input variability on EEG-BOLD correlation (cf. case 1 in Figure 5). (**A**) K-means clusters (same as in Figure 2**B**) within the input space *I*_lat_ = 4.5, *I*_mod_ = [0–25], and *I*_tha_ = [0–5]. The effect of external input variability (i.e., common driving inputs to specific populations) on alpha-BOLD and gamma-BOLD correlations is examined by incrementally increasing the input ranges (indicated by the black rectangles). The progression starts from the constant input condition (*I*_mod_ = 1 and *I*_tha_ = 1) to the varying input condition (*I*_mod_ = [0–2] and *I*_tha_ = [0–2]), as shown in Figure 5, case 1. (**B**) Histograms of alpha-BOLD (blue) and gamma-BOLD (orange) correlations across 100 runs for each step of the varying input range. The gradual increase in mean alpha-BOLD *r*_*α*_ correlation and mean gamma-BOLD correlation *r*_*γ*_ is evident, so only the first few steps and the final step are displayed. (**C**) Example time series from a single 10-minute run under the constant input condition (left) and the varying input condition (right). (**D**) Mean correlation of the firing rates of 17 populations under the constant input condition (left), the varying input condition (middle), and the difference between the two conditions (right). The populational coherence (Coh) is calculated as the mean absolute values of the correlation matrix (excluding the diagonal part).

**Figure S9.**
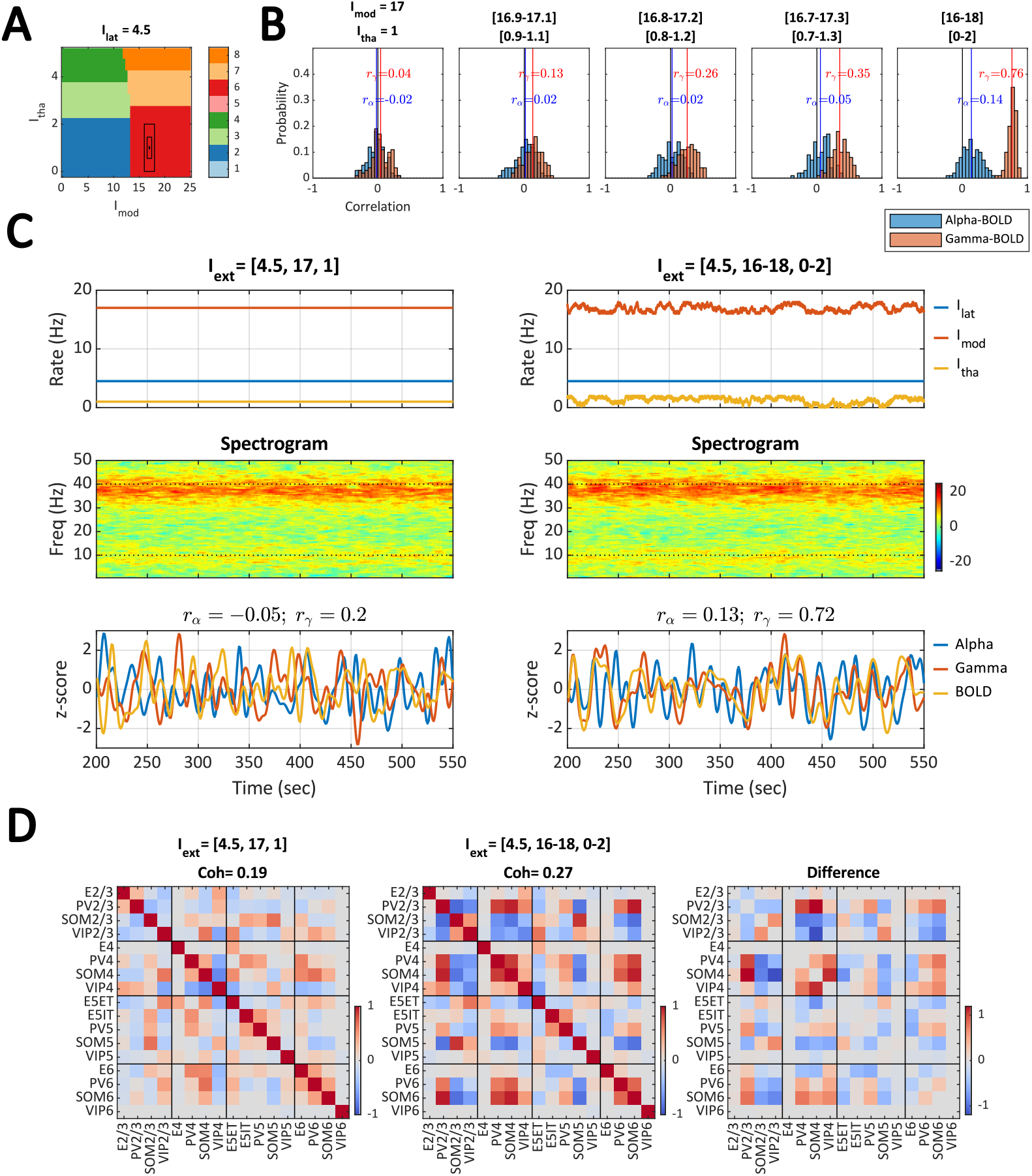
Effect of external input variability on EEG-BOLD correlation (cf. case 3 in Figure 5). (**A**) K-means clusters (same as in Figure 2**B**) within the input space *I*_lat_ = 4.5, *I*_mod_ = [0–25], and *I*_tha_ = [0–5]. The effect of external input variability (i.e., common driving inputs to specific populations) on alpha-BOLD and gamma-BOLD correlations is examined by incrementally increasing the input ranges (indicated by the black rectangles). The progression starts from the constant input condition (*I*_mod_ = 17 and *I*_tha_ = 1) to the varying input condition (*I*_mod_ = [16–18] and *I*_tha_ = [0–2]), as shown in Figure 5, case 3. (**B**) Histograms of alpha-BOLD (blue) and gamma-BOLD (orange) correlations across 100 runs for each step of the varying input range. The gradual increase in mean alpha-BOLD correlation *r*_*α*_ and mean gamma-BOLD correlation *r*_*γ*_ is evident, so only the first few steps and the final step are displayed. (**C**) Example time series from a single 10-minute run under the constant input condition (left) and the varying input condition (right). (**D**) Mean correlation of the firing rates of 17 populations under the constant input condition (left), the varying input condition (middle), and the difference between the two conditions (right). The populational coherence (Coh) is calculated as the mean absolute values of the correlation matrix (excluding the diagonal part).

**Figure S10.**
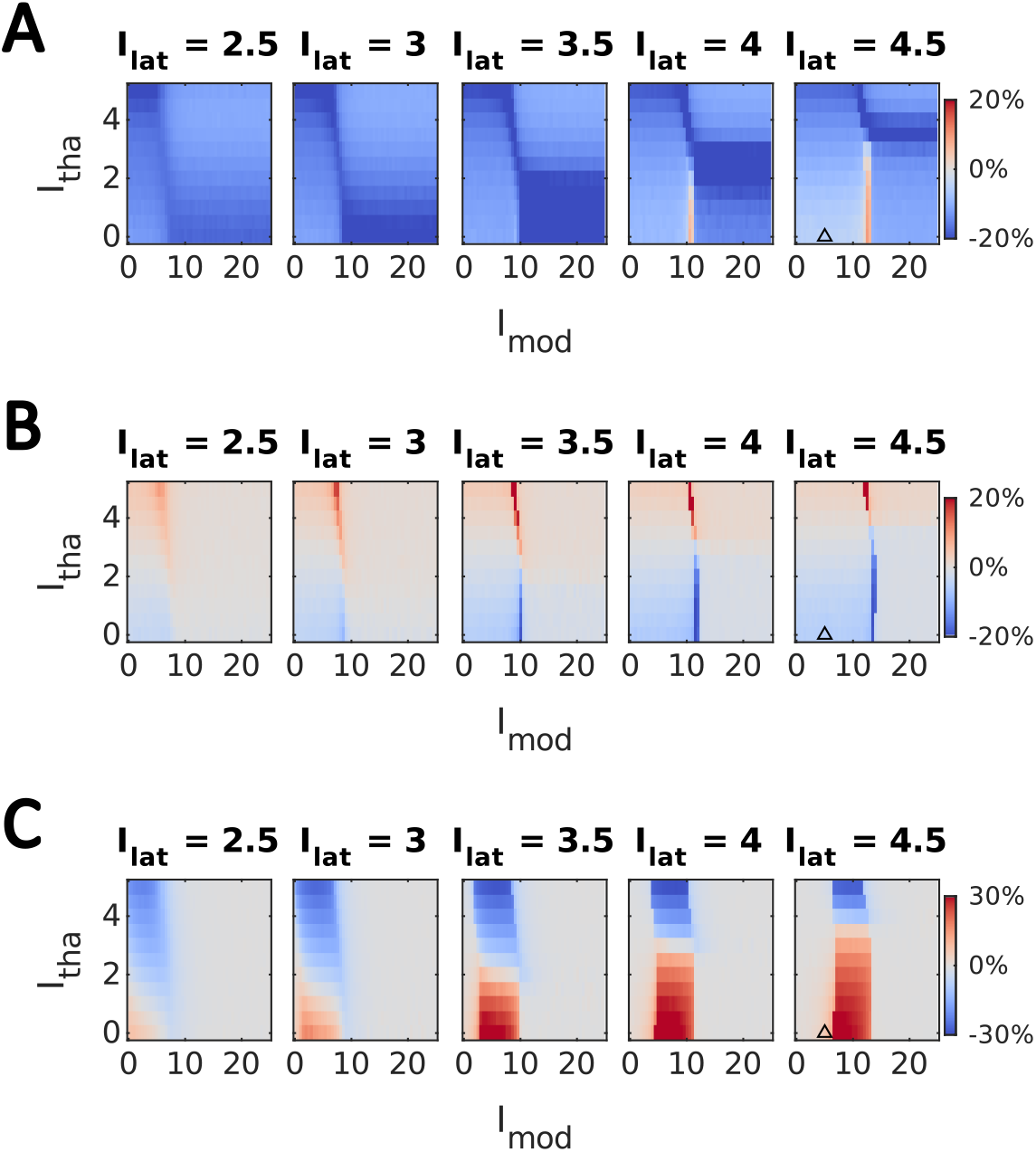
Polarity of BOLD responses to selective stimulation of interneuron subtypes. Relative changes in simulated BOLD signals (color-coded as percentages) are shown across the input parameter space when an additional step input (duration = 20 sec) is applied selectively to one of three interneuron populations: (**A**) PV2/3, (**B**) SOM2/3, or (**C**) VIP2/3. The triangle marks the input configuration [*I*_lat_, *I*_mod_, *I*_tha_] = [4.5, 5, 0], which is used as the illustrative example in Figure 9.

## Notes

### Competing Interest Statement

The authors have declared no competing interest.

### Summary of Updates

Added one more subsection, "Circuit-level effect on BOLD response to optogenetic stimulation."

## Bibliography

Adesnik, H., Bruns, W., Taniguchi, H., Huang, Z. J., & Scanziani, M. (2012). A neural circuit for spatial summation in visual cortex. Nature, 490(7419), 226–231.

Barry, R. J., Clarke, A. R., Johnstone, S. J., Magee, C. A., & Rushby, J. A. (2007). Eeg differences between eyes-closed and eyes-open resting conditions. Clinical neurophysiology, 118(12), 2765–2773.

Bastos, G., Holmes, J. T., Ross, J. M., Rader, A. M., Gallimore, C. G., Wargo, J. A., Peterka, D. S., & Hamm, J. P. (2023). Top-down input modulates visual context processing through an interneuron-specific circuit. Cell reports, 42(9).

Berger, H. (1929). Über das elektroenkephalogramm des menschen. Archiv für psychiatrie und nervenkrankheiten, 87 (1), 527–570.

Biasiucci, A., Franceschiello, B., & Murray, M. M. (2019). Electroencephalography. Current Biology, 29(3), R80–R85.

Buzsáki, G., Anastassiou, C. A., & Koch, C. (2012). The origin of extracellular fields and currents—eeg, ecog, lfp and spikes. Nature reviews neuroscience, 13(6), 407–420.

Campagnola, L., Seeman, S. C., Chartrand, T., Kim, L., Hoggarth, A., Gamlin, C., Ito, S., Trinh, J., Davoudian, P., Radaelli, C., et al. (2022). Local connectivity and synaptic dynamics in mouse and human neocortex. Science, 375(6585), eabj5861.

Cauli, B., Tong, X.-K., Rancillac, A., Serluca, N., Lambolez, B., Rossier, J., & Hamel, E. (2004). Cortical gaba interneurons in neurovascular coupling: Relays for subcortical vasoactive pathways. Journal of Neuroscience, 24(41), 8940–8949.

Chen, G., Zhang, Y., Li, X., Zhao, X., Ye, Q., Lin, Y., Tao, H. W., Rasch, M. J., & Zhang, X. (2017). Distinct inhibitory circuits orchestrate cortical beta and gamma band oscillations. Neuron, 96(6), 1403–1418.

Chien, V. S., Wang, P., Maess, B., Fishman, Y., & Knösche, T. R. (2023). Laminar neural dynamics of auditory evoked responses: Computational modeling of local field potentials in auditory cortex of non-human primates. NeuroImage, 281, 120364.

Clayton, M. S., Yeung, N., & Cohen Kadosh, R. (2018). The many characters of visual alpha oscillations. European Journal of Neuroscience, 48(7), 2498–2508.

Cona, G., Chiossi, F., Di Tomasso, S., Pellegrino, G., Piccione, F., Bisiacchi, P., & Arcara, G. (2020). Theta and alpha oscillations as signatures of internal and external attention to delayed intentions: A magnetoencephalography (meg) study. NeuroImage, 205, 116295.

Domhof, J. W., & Tiesinga, P. H. (2021). Flexible frequency switching in adult mouse visual cortex is mediated by competition between parvalbumin and somatostatin expressing interneurons. Neural Computation, 33(4), 926–966.

Ekstrom, A. (2010). How and when the fmri bold signal relates to underlying neural activity: The danger in dissociation. Brain research reviews, 62(2), 233–244.

Fang, R., Xia, C., Close, J. L., Zhang, M., He, J., Huang, Z., Halpern, A. R., Long, B., Miller, J. A., Lein, E. S., et al. (2022). Conservation and divergence of cortical cell organization in human and mouse revealed by merfish. Science, 377 (6601), 56–62.

Friston, K. J., Harrison, L., & Penny, W. (2003). Dynamic causal modelling. Neuroimage, 19(4), 1273–1302.

Friston, K. J., Mechelli, A., Turner, R., & Price, C. J. (2000). Nonlinear responses in fmri: The balloon model, volterra kernels, and other hemodynamics. NeuroImage, 12(4), 466– 477.

Goldman, R. I., Stern, J. M., Engel Jr, J., & Cohen, M. S. (2002). Simultaneous eeg and fmri of the alpha rhythm. Neuroreport, 13(18), 2487–2492.

Gonçalves, S. I., De Munck, J. C., Pouwels, P. J., Schoonhoven, R., Kuijer, J. P., Maurits, N. M., Hoogduin, J. M., Van Someren, E. J., Heethaar, R. M., & Da Silva, F. L. (2006). Correlating the alpha rhythm to bold using simultaneous eeg/fmri: Inter-subject variability. Neuroimage, 30(1), 203–213.

Gonchar, Y., Wang, Q., & Burkhalter, A. H. (2008). Multiple distinct subtypes of gabaergic neurons in mouse visual cortex identified by triple immunostaining. Frontiers in neuroanatomy, 2, 93.

Grinvald, A., Lieke, E. E., Frostig, R. D., & Hildesheim, R. (1994). Cortical point-spread function and long-range lateral interactions revealed by real-time optical imaging of macaque monkey primary visual cortex. Journal of Neuroscience, 14(5), 2545– 2568.

Hahn, G., Kumar, A., Schmidt, H., Knösche, T. R., & Deco, G. (2022). Rate and oscillatory switching dynamics of a multilayer visual microcircuit model. Elife, 11, e77594.

Halgren, M., Ulbert, I., Bastuji, H., Fabó, D., Erő ss, L., Rey, M., Devinsky, O., Doyle, W. K., Mak-McCully, R., Halgren, E., et al. (2019). The generation and propagation of the human alpha rhythm. Proceedings of the National Academy of Sciences, 116(47), 23772–23782.

Hertäg, L., & Sprekeler, H. (2019). Amplifying the redistribution of somato-dendritic inhibition by the interplay of three interneuron types. PLoS computational biology, 15(5), e1006999.

Howarth, C., Gleeson, P., & Attwell, D. (2012). Updated energy budgets for neural computation in the neocortex and cerebellum. Journal of Cerebral Blood Flow & Metabolism, 32(7), 1222–1232.

Howarth, C., Mishra, A., & Hall, C. N. (2021). More than just summed neuronal activity: How multiple cell types shape the bold response. Philosophical Transactions of the Royal Society B, 376(1815), 20190630.

Jansen, B. H., & Rit, V. G. (1995). Electroencephalogram and visual evoked potential generation in a mathematical model of coupled cortical columns. Biological cybernetics, 73(4), 357–366.

Ji, X.-y., Zingg, B., Mesik, L., Xiao, Z., Zhang, L. I., & Tao, H. W. (2016). Thalamocortical innervation pattern in mouse auditory and visual cortex: Laminar and cell-type specificity. Cerebral cortex, 26(6), 2612–2625.

Jiang, H.-J., Qi, G., Duarte, R., Feldmeyer, D., & van Albada, S. J. (2024). A layered microcircuit model of somatosensory cortex with three interneuron types and cell-type-specific short-term plasticity. Cerebral Cortex, 34(9), bhae378.

Jolivet, R., Coggan, J. S., Allaman, I., & Magistretti, P. J. (2015). Multi-timescale modeling of activity-dependent metabolic coupling in the neuron-glia-vasculature ensemble. PLoS computational biology, 11(2), e1004036.

Keller, A. S., Payne, L., & Sekuler, R. (2017). Characterizing the roles of alpha and theta oscillations in multisensory attention. Neuropsychologia, 99, 48–63.

Kloc, M., & Maffei, A. (2014). Target-specific properties of thalamocortical synapses onto layer 4 of mouse primary visual cortex. Journal of Neuroscience, 34(46), 15455–15465.

Krawchuk, M. B., Ruff, C. F., Yang, X., Ross, S. E., & Vazquez, A. L. (2020). Optogenetic assessment of vip, pv, som and nos inhibitory neuron activity and cerebral blood flow regulation in mouse somato-sensory cortex. Journal of Cerebral Blood Flow & Metabolism, 40(7), 1427–1440.

Laufs, H., Kleinschmidt, A., Beyerle, A., Eger, E., Salek-Haddadi, A., Preibisch, C., & Krakow, K. (2003). Eeg-correlated fmri of human alpha activity. Neuroimage, 19(4), 1463– 1476.

Laufs, H., Krakow, K., Sterzer, P., Eger, E., Beyerle, A., Salek-Haddadi, A., & Kleinschmidt, A. (2003). Electroencephalographic signatures of attentional and cognitive default modes in spontaneous brain activity fluctuations at rest. Proceedings of the national academy of sciences, 100(19), 11053–11058.

Lee, B., Shin, D., Gross, S. P., & Cho, K.-H. (2018). Combined positive and negative feedback allows modulation of neuronal oscillation frequency during sensory processing. Cell Reports, 25(6), 1548–1560.

Lee, L., Boorman, L., Glendenning, E., Christmas, C., Sharp, P., Redgrave, P., Shabir, O., Bracci, E., Berwick, J., & Howarth, C. (2020). Key aspects of neurovascular control mediated by specific populations of inhibitory cortical interneurons. Cerebral Cortex, 30(4), 2452–2464.

Logothetis, N. K., Pauls, J., Augath, M., Trinath, T., & Oeltermann, A. (2001). Neurophysiological investigation of the basis of the fmri signal. nature, 412(6843), 150–157.

Magosso, E., De Crescenzio, F., Ricci, G., Piastra, S., & Ursino, M. (2019). Eeg alpha power is modulated by attentional changes during cognitive tasks and virtual reality immersion. Computational intelligence and neuroscience, 2019(1), 7051079.

Magri, C., Schridde, U., Murayama, Y., Panzeri, S., & Logothetis, N. K. (2012). The amplitude and timing of the bold signal reflects the relationship between local field potential power at different frequencies. Journal of Neuroscience, 32(4), 1395–1407.

Missonnier, P., Deiber, M.-P., Gold, G., Millet, P., Gex-Fabry Pun, M., Fazio-Costa, L., Giannakopoulos, P., & Ibáñez, V. (2006). Frontal theta event-related synchronization: Comparison of directed attention and working memory load effects. Journal of neural transmission, 113, 1477–1486.

Moon, H. S., Jiang, H., Vo, T. T., Jung, W. B., Vazquez, A. L., & Kim, S.-G. (2021). Contribution of excitatory and inhibitory neuronal activity to bold fmri. Cerebral Cortex, 31(9), 4053–4067.

Moosmann, M., Ritter, P., Krastel, I., Brink, A., Thees, S., Blankenburg, F., Taskin, B., Obrig, H., & Villringer, A. (2003). Correlates of alpha rhythm in functional magnetic resonance imaging and near infrared spectroscopy. Neuroimage, 20(1), 145–158.

Muthukumaraswamy, S. D., & Singh, K. D. (2013). Visual gamma oscillations: The effects of stimulus type, visual field coverage and stimulus motion on meg and eeg recordings. Neuroimage, 69, 223–230.

Næss, S., Halnes, G., Hagen, E., Hagler Jr, D. J., Dale, A. M., Einevoll, G. T., & Ness, T. V. (2021). Biophysically detailed forward modeling of the neural origin of eeg and meg signals. NeuroImage, 225, 117467.

Nakagawa, T. T., Woolrich, M., Luckhoo, H., Joensson, M., Mohseni, H., Kringelbach, M. L., Jirsa, V., & Deco, G. (2014). How delays matter in an oscillatory whole-brain spiking-neuron network model for meg alpha-rhythms at rest. NeuroImage, 87, 383–394.

Niessing, J., Ebisch, B., Schmidt, K. E., Niessing, M., Singer, W., & Galuske, R. A. (2005). Hemodynamic signals correlate tightly with synchronized gamma oscillations. science, 309(5736), 948–951.

Onorato, I., Tzanou, A., Schneider, M., Uran, C., Broggini, A. C., & Vinck, M. (2025). Distinct roles of pv and sst interneurons in visually induced gamma oscillations. Cell Reports, 44(3).

Pang, J. C., & Robinson, P. A. (2018). Neural mechanisms of the eeg alpha-bold anticorrelation. Neuroimage, 181, 461–470.

Sanchez-Todo, R., Bastos, A. M., Lopez-Sola, E., Mercadal, B., Santarnecchi, E., Miller, E. K., Deco, G., & Ruffini, G. (2023). A physical neural mass model framework for the analysis of oscillatory generators from laminar electrophysiological recordings. NeuroImage, 270, 119938.

Schirner, M., McIntosh, A. R., Jirsa, V., Deco, G., & Ritter, P. (2018). Inferring multi-scale neural mechanisms with brain network modelling. elife, 7, e28927.

Shu, I.-W., Lin, Y., Granholm, E. L., & Singh, F. (2024). A focused review of gamma neuromodulation as a therapeutic target in alzheimer’s spectrum disorders. Journal of psychiatry and brain science, 9(1).

Spiegler, A., Kiebel, S. J., Atay, F. M., & Knösche, T. R. (2010). Bifurcation analysis of neural mass models: Impact of extrinsic inputs and dendritic time constants. NeuroImage, 52(3), 1041–1058.

Uhlirova, H., Kılıç, K., Tian, P., Thunemann, M., Desjardins, M., Saisan, P. A., Sakadžić, S., Ness, T. V., Mateo, C., Cheng, Q., et al. (2016). Cell type specificity of neurovascular coupling in cerebral cortex. elife, 5, e14315.

Vazquez, A. L., Fukuda, M., & Kim, S.-G. (2018). Inhibitory neuron activity contributions to hemodynamic responses and metabolic load examined using an inhibitory optogenetic mouse model. Cerebral Cortex, 28(11), 4105–4119.

Veit, J., Hakim, R., Jadi, M. P., Sejnowski, T. J., & Adesnik, H. (2017). Cortical gamma band synchronization through somatostatin interneurons. Nature neuroscience, 20(7), 951–959.

Vierling-Claassen, D., Cardin, J. A., Moore, C. I., & Jones, S. R. (2010). Computational modeling of distinct neocortical oscillations driven by cell-type selective optogenetic drive: Separable resonant circuits controlled by low-threshold spiking and fast-spiking interneurons. Frontiers in human neuroscience, 4, 198.

Wang, P., & Knösche, T. R. (2013). A realistic neural mass model of the cortex with laminar-specific connections and synaptic plasticity–evaluation with auditory habituation. PloS one, 8(10), e77876.

Whittington, M. A., & Traub, R. D. (2003). Interneuron diversity series: Inhibitory interneurons and network oscillations in vitro. Trends in neurosciences, 26(12), 676–682.

Yavorska, I., & Wehr, M. (2016). Somatostatin-expressing inhibitory interneurons in cortical circuits. Frontiers in neural circuits, 10, 76.

## Supplementary Bibliography

Delorme, A., & Makeig, S. (2004). Eeglab: An open source toolbox for analysis of single-trial eeg dynamics including independent component analysis. Journal of neuroscience methods, 134(1), 9–21.

Glover, G. H., Li, T.-Q., & Ress, D. (2000). Image-based method for retrospective correction of physiological motion effects in fmri: Retroicor. Magnetic Resonance in Medicine: An Official Journal of the International Society for Magnetic Resonance in Medicine, 44(1), 162–167.

Jiricek, S., Koudelka, V., Mantini, D., Marecek, R., & Hlinka, J. (2025). Spatial (mis)match between eeg and fmri signal patterns revealed by spatio-spectral source-space eeg decomposition. Frontiers in Neuroscience, Volume 19 - 2025. 10.3389/fnins.2025.1549172

Liu, Q., Farahibozorg, S., Porcaro, C., Wenderoth, N., & Mantini, D. (2017). Detecting large-scale networks in the human brain using high-density electroencephalography. Human brain mapping, 38(9), 4631–4643.

Liu, Q., Ganzetti, M., Wenderoth, N., & Mantini, D. (2018). Detecting large-scale brain networks using eeg: Impact of electrode density, head modeling and source localization. Frontiers in neuroinformatics, 12, 4.

Marino, M., Arcara, G., Porcaro, C., & Mantini, D. (2019). Hemodynamic correlates of electrophysiological activity in the default mode network. Frontiers in neuroscience, 13, 1060.

Marino, M., Liu, Q., Koudelka, V., Porcaro, C., Hlinka, J., Wenderoth, N., & Mantini, D. (2018). Adaptive optimal basis set for bcg artifact removal in simultaneous eeg-fmri. Scientific reports, 8(1), 1–11

Niazy, R. K., Beckmann, C. F., Iannetti, G. D., Brady, J. M., & Smith, S. M. (2005). Removal of fmri environment artifacts from eeg data using optimal basis sets. Neuroimage, 28(3), 720–737.

Oostenveld, R., Fries, P., Maris, E., & Schoffelen, J.-M. (2011). Fieldtrip: Open source software for advanced analysis of meg, eeg, and invasive electrophysiological data. Computational intelligence and neuroscience, 2011.

Pascual-Marqui, R. D., Lehmann, D., Koukkou, M., Kochi, K., Anderer, P., Saletu, B., Tanaka, H., Hirata, K., John, E. R., Prichep, L., et al. (2011). Assessing interactions in the brain with exact low-resolution electromagnetic tomography. Philosophical Transactions of the Royal Society A: Mathematical, Physical and Engineering Sciences, 369(1952), 3768–3784.

Penny, W. D., Friston, K. J., Ashburner, J. T., Kiebel, S. J., & Nichols, T. E. (2011). Statistical parametric mapping: The analysis of functional brain images. Elsevier.

Taberna, G. A., Samogin, J., & Mantini, D. (2021). Automated head tissue modelling based on structural magnetic resonance images for electroencephalographic source reconstruction. Neuroinformatics, 1–12.

Xia, M., Wang, J., & He, Y. (2013). Brainnet viewer: A network visualization tool for human brain connectomics. PloS one, 8(7), e68910.

